# Super-resolution Imaging Reveals 3D Structure and Organizing Mechanism of Accessible Chromatin

**DOI:** 10.1101/678649

**Authors:** Liangqi Xie, Peng Dong, Yifeng Qi, Margherita De Marzio, Xingqi Chen, Sambashiva Banala, Wesley R. Legant, Brian P. English, Anders S. Hansen, Anton Schulmann, Luke D. Lavis, Eric Betzig, Rafael Casellas, Howard Y. Chang, Bin Zhang, Robert Tjian, Zhe Liu

## Abstract

*Access* to *cis*-regulatory elements packaged in chromatin is essential for directing gene expression and cell viability. Here, we report a super-resolution imaging strategy, 3D ATAC-PALM, that enables direct visualization of the entire accessible genome. We found that active chromosomal segments are organized into spatially-segregated accessible chromatin domains (ACDs). Rapid depletion of CTCF or Cohesin (RAD21 subunit) induced enhanced ACD clustering, reduced physical separation between intrachromosomal ACDs, and differentially regulated ACD compaction. Experimental perturbations and polymer modeling suggest that dynamic protein-protein and protein-DNA interactions within ACDs couple with loop extrusion to organize ACD topology. Dysorganization of ACDs upon CTCF or Cohesin loss alters transcription factor binding and target search dynamics in living cells. These results uncover fundamental mechanisms underpinning the formation of 3D genome architecture and its pivotal function in transcriptional regulation.

## INTRODUCTION

The rapid development of genomic methods derived from chromosome conformation capture (3C) assays in the past decade has provided significant measurements to probe chromatin folding and genome organization with rich sequence information (Dekker and Mirny, 2016; Dekker et al., 2002, 2013; Wit and Laat, 2012). The general model emerging from such studies is that the mammalian genome is organized into distinct architectural scales including compartments (Lieberman-Aiden et al., 2009), topologically associated domains (TADs) (Dixon et al., 2012; Nora et al., 2012; Sexton et al., 2012), and loop domains (Rao et al., 2014). Two key protein components (*e.g.* CTCF and Cohesin) were identified that cooperate to build TADs and loop domains by the loop extrusion mechanism (Alipour and Marko, 2012; Fudenberg et al., 2016; Sanborn et al., 2015). Single-cell Hi-C and high-throughput DNA fluorescence *in situ* hybridization (FISH) have also revealed substantial structural variability in the organization of TADs and loops within individual cells (Bintu et al., 2018; Finn et al., 2019; Flyamer et al., 2017; Stevens et al., 2017). These findings suggest that reconstructing three-dimensional (3D) genome organization based on ensemble 3C-based measurements is limited by the inherent stochasticity of loop formation and temporal fluctuations in the state of chromatin.

Of the ~6 billion base pairs of DNA in the diploid human genome, only a small fraction (~2%) is accessible for binding by transcription factors (TFs), which decode essential genetic information to instruct precise spatiotemporal gene expression programs (Levine et al., 2014). The linear distribution of accessible chromatin (*e.g.* enhancers, promoters and insulators) has been extensively mapped by DNase I digestion (Boyle et al., 2008; Wu, 1980) or by assay for transposase-accessible chromatin with high-throughput sequencing (ATAC-seq) (Buenrostro et al., 2013). Recently, an ATAC based imaging method (ATAC-See) has been developed, revealing cell-type specific accessible chromatin organization in single cells (Chen et al., 2016). However, because of the diffraction-limited resolution, this technique was unable to detect finer chromatin structures inside the compact nuclei where relevant structure and organizing mechanism may be revealed.

To explore genome organization at super resolution, in particular at gene regulatory elements, we devised an imaging strategy to visualize the 3D structure of the entire accessible genome at nanometer-scale. Specifically, we used the Tn5 transposase to insert photo-activatable fluorescent DNA probes into the accessible genome of mouse embryonic stem cells (ESCs) (Chen et al., 2016; Grimm et al., 2016) (**Fig. 1A, Fig. S1A-G**). To achieve whole nucleus 3D imaging beyond the diffraction limit, we applied photoactivated localization microscopy (PALM) (Betzig et al., 2006). We took advantage of the ultra-thin axial sectioning of the Lattice Light Sheet microscope (LLSM) (Chen et al., 2014a; Legant et al., 2016) and engineered a depth-sensitive point spread function (PSF) by astigmatism in order to improve axial localization precision (< 50 nm) (Huang et al., 2008; Legant et al., 2016) (**Fig. 1B**). This strategy minimized out-of-focus photobleaching and maximized the photon budget for accurate single molecule localization throughout the nucleus. We named this strategy “3D ATAC-PALM”.

**Figure 1.**
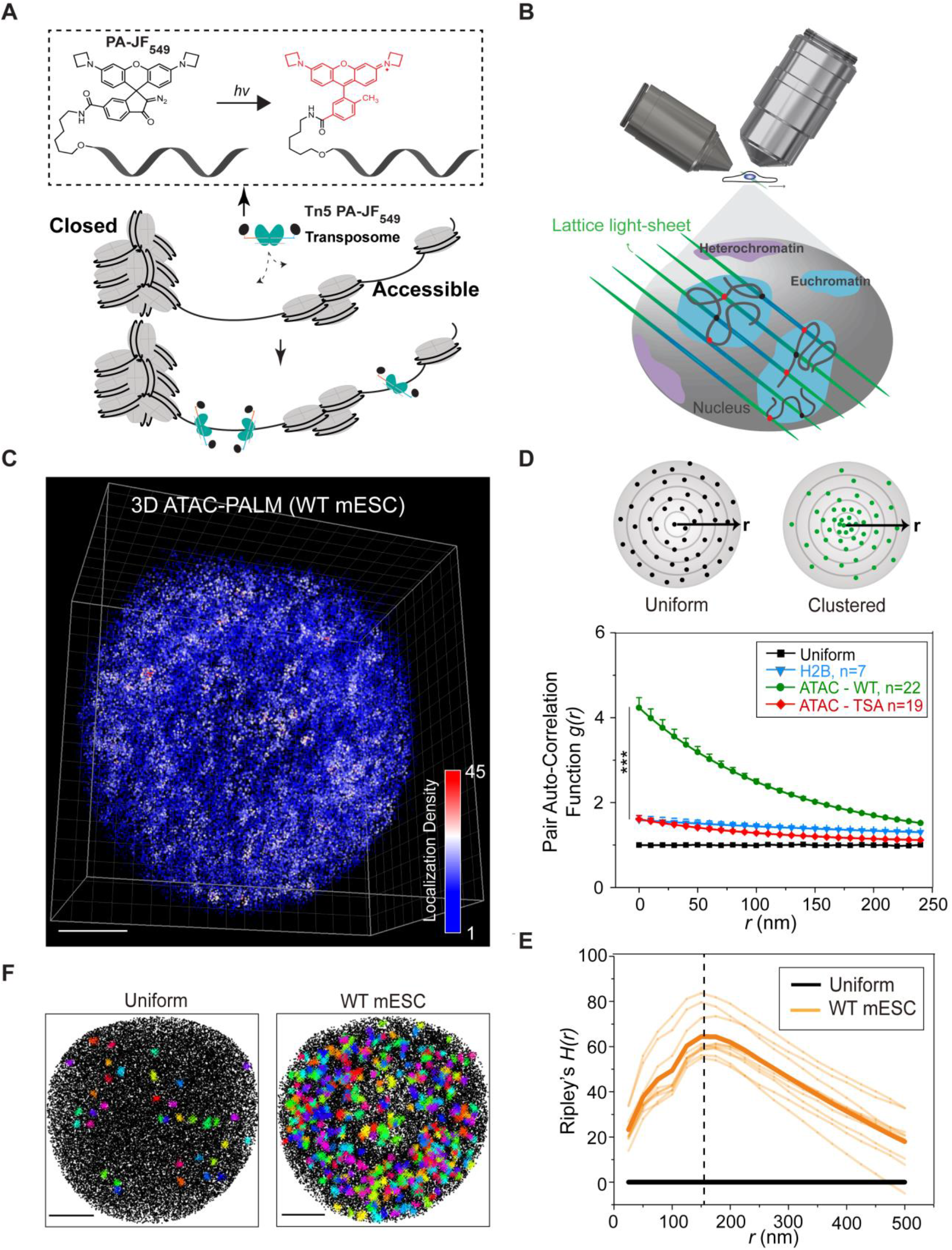
Visualizing the accessible genome topology by 3D ATAC-PALM microscopy. **(A)** Schematics of the 3D ATAC-PALM microscopy labeling and imaging strategy. Photoactivatable Janelia Fluor 549 (PA-JF_549_) was conjugated to DNA oligo containing the mosaic ends (ME) of the Tn5 transposon and reconstituted with Tn5 transposase (cyan) to form the active transposome complex *in vitro.* The cells were fixed, permeabilized, and accessible sites in the genome were selectively labeled by the Tn5 PA-JF_549_ transposome. Photoactivation (hv, 405 nm laser) of the nonfluorescent PA-JF_549_ yields highly fluorescent methyl-substituted JF_549_. **(B)** Tn5 PA-JF_549_ transposome treated cells were mounted onto the Lattice Light Sheet microscope for 3D ATAC-PALM imaging. **(C)** 3D illustration of accessible chromatin localizations in wild-type (WT) mouse ESCs. The color-coded localization density was calculated with a canopy radius of 250nm. Scale bar: 2 μm. **(D)** Global pair auto-correlation function *g(r)* analysis of ATAC-PALM localizations for WT and TSA treated ESCs. The top panel shows a simplified 2D scheme for uniform (black dots, top left) or clustered (green dots, top right) distribution of localizations. *g(r)* represents the pair auto-correlation function of distance *r* calculated from a given origin point (the center of all circles) within the space. Our analysis was performed based on 3D coordinates. The *g(r)* for 3D H2B-HaloTag PALM localizations was computed as a control. The error bars represent standard error (SE) of the mean. The non-parametric Mann-Whitney U test was used to compare the clustering amplitude *A* (equals to *g(0))* in different groups. *g(r)* was plotted from the fitted exponential decay function. **(E)** Ripley’s H function *H(r)* of the ACDs from WT mESCs. The black line indicates uniform distribution and the light-yellow lines represent the *H(r)* from individual WT mESCs (n=10) and the thick yellow line indicate their mean. The vertical dashed line represents the peak radius position around 150 nm. **(F)** Identification of ACDs by DBSCAN algorithm. Both panels show the Z-projection of all the localizations (black dots) and identified clusters (colored crosses). Left panel, DBSCAN only detected 36 clusters in uniformly sampled data points with the same average density as in the right panel. Right panel, DBSCAN identified 402 clusters in WT ESCs. Scale bar, 2 μm. *, p < 0.05; *, p < 0.01; ***, p < 0.001; n.s., not significant; throughout the publication unless specified.

### The accessible genome is organized into spatially-segregated ACDs

After nucleus segmentation and 3D ATAC-PALM reconstruction, one consistent feature observed in different cell types and across distinct cell cycle stages is that the accessible genome is organized into spatially-segregated 3D clusters hereafter called <u>a</u>ccessible <u>c</u>hromatin <u>d</u>omains (ACDs) (**Fig. 1C, Fig. S2, Fig. S3, C-D, Movie S1**). To quantify the extent of accessible chromatin clustering, we used the pair auto-correlation function *g(r)* which increases in value as localizations become more clustered (**Fig. 1D; Equation S5–7**) (Peebles, 1973). The mean radius of ACDs in ESCs is ~ 150nm, as determined by Ripley’s *H(r)* analysis (**Equation S8–10**) (**Fig. 1E**). The number of ACDs in ESCs is ~ 11 fold larger than that for uniformly sampled localizations in the same volume, as determined by DBSCAN (Density-Based Spatial Clustering of Applications with Noise) analysis (**Fig. 1F**) (Ester, M., Kriegel, H. P., Sander, J., & Xu, 1996). Importantly, ATAC-PALM localizations are significantly more clustered than histone H2B PALM localizations (**Fig. 1D**), excluding the possibility that over-counting of blinking molecules could account for 3D clustering of accessible chromatin. To test whether we can perturb the organization of ACDs, we induced chromatin hyperacetylation by the HDAC inhibitor trichostatin A (TSA) (Kieffer-Kwon et al., 2017). TSA treatment markedly reduced *g(r)* (**Fig. 1D**), suggesting less clustering and a more homogenous distribution of accessible chromatin in the nucleus (*Fig. S3, A and B*). Thus, 3D ATAC-PALM can detect and quantify 3D organizational changes of accessible chromatin.

To further verify whether 3D ATAC-PALM targets accessible chromatin, we imaged ESCs stably expressing heterochromatin protein 1 fused with GFP (HP1-GFP) and found that ACDs and HP1-GFP labeled heterochromatic regions are mutually exclusive, organized into distinct salt-and-pepper spatial patterns in the nucleus (*Fig. S4A, Movie S2*). In addition, by coupling 3D ATAC-PALM with 3D Oligopaint FISH (Beliveau et al., 2012, 2015), we found that ACDs are spatially co-localized with well-defined active (ATAC-rich) chromosomal segments (*Fig. S4, B-D, Movie S3-S4*) but not with inactive (ATAC-poor) ones (*Fig. S4, E and F*). Taken together, these results demonstrated that ACDs represent structures of active, accessible chromatin that are spatially segregated from heterochromatin.

### Spatial re-organization of ACDs upon acute loss of CTCF or Cohesin

To investigate mechanisms controlling the spatial organization of ACDs, we focused on CTCF and Cohesin, which are thought to regulate genome organization through loop extrusion (Alipour and Marko, 2012; Fudenberg et al., 2016; Sanborn et al., 2015). To perturb CTCF- and Cohesin-mediated functions with minimal secondary effects, we employed the auxin-inducible degron (AID) system (Nishimura et al., 2009) and tagged endogenous CTCF or Cohesin subunit RAD21 with a HaloTag-miniAID (mAID) in a mouse ESC line stably expressing the rice F-box protein TIR1 (**Fig. 2A**). The HaloTag-mAID was fused to the N-terminus of CTCF to avoid previously reported problems associated with C-terminal tagging (*e.g.* auxin independent degradation and interference with CTCF function) (Wutz et al., 2017). We found that HaloTag-mAID labeling of CTCF (N-terminal) or RAD21(C-terminal) influenced neither basal protein levels (**Fig. S5, A and B**) nor cell proliferation before auxin treatment (**Fig. S6, A, B and D**), consistent with our previous findings that tagging CTCF or RAD21 at these positions does not interfere with protein function (Hansen et al., 2017). Within a few hours of auxin treatment, CTCF or RAD21 expression was undetectable by western blot, single cell fluorescence imaging or flow cytometry (> 99% depletion for CTCF) (**Fig. 2A, Fig. S5**). The depletion was reversible, as CTCF or RAD21 protein levels quickly recovered after auxin washout (**Fig. S5, A and D**). We found that the acute loss of CTCF (up to 12 hours) or RAD21 (up to 6 hours) did not cause noticeable changes in proliferation, cell cycle phasing or expression of pluripotency markers (**Fig. 2A, Fig. S5, A and C, Fig. S6, A and B**). However, prolonged CTCF (> 48 hours) or RAD21 (> 24 hours) loss did compromise proliferation, colony formation and survival of ESCs (**Fig. S6**).

**Figure 2.**
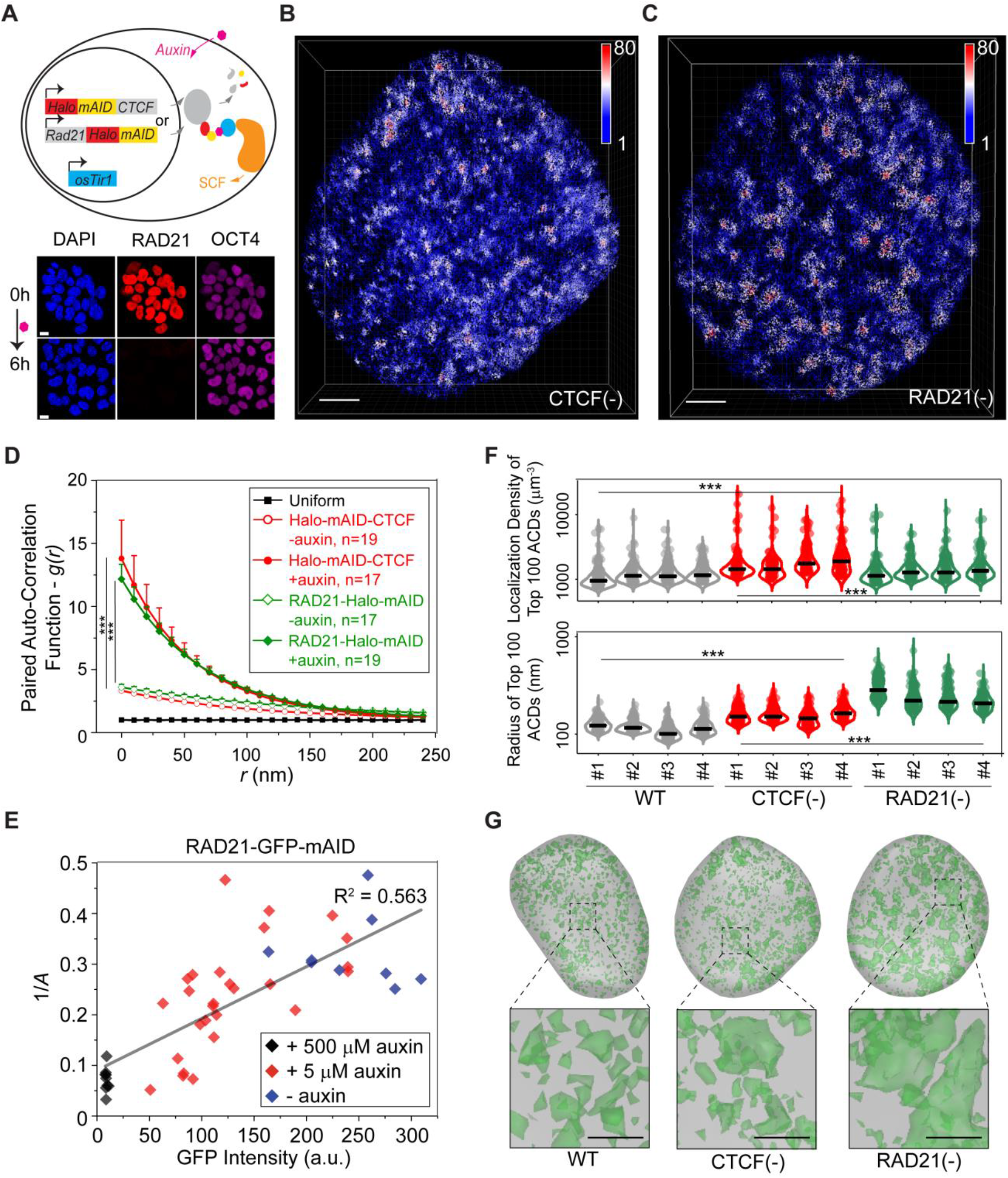
Structural variation of accessible chromatin upon acute loss of CTCF or Cohesin. (**A**) Acute depletion of CTCF and Cohesin in mESCs by the auxin-induced protein depletion system. The mini auxin-inducible degron (mAID)-HaloTag (halo) was bi-allelically knocked into N-terminus of *CTCF* or C-terminus of *Rad21* (Cohesin subunit) by CRISPR/Cas9 genome editing technology (upper panel) in a mouse ES line stably expressing the plant derived E3 ligase adaptor protein osTir1. Adding plant derived hormone analogue (auxin) triggers rapid degradation of target protein RAD21 or CTCF (See also Figure S5A-B) as revealed by single cell immunofluorescence (lower panel). OCT4 immunostaining was used as a control. Scale bar, 5 μm. (**B** and **C**) Single-cell 3D illustration of ATAC-PALM localizations upon CTCF (**B**) and RAD21 (**C**) depletion. The color bar indicates localization density calculated by using a canopy radius of 250 nm. See the 3D rotatory presentation in **Movie S5**. Scale bar, 2 μm. (**D**) CTCF and RAD21 depletion promotes global accessible chromatin clustering. The error bar represents standard error (SE) of the mean and Mann-Whitney U test was applied for comparing data points at *g(0)*. (**E**) The dose-dependent effect of Cohesin depletion on global accessible chromatin clustering revealed by the inverse relationship between clustering amplitude (*A*) and residual RAD21 levels measured by RAD21-GFP-mAID fluorescence intensities (arbitrary fluorescent units). Specifically, two different auxin concentrations (5 μM and 500 μM) were used to generate a gradient of RAD21 level in single cells. (**F**) Acute CTCF or RAD21 depletion affects ACD formation through distinct mechanisms. The upper panel shows the violin plot of localization density of top 100 rank ordered ACDs among 4 individual cells for each condition. The lower panel shows the violin plot of normalized radius of top 100 ranked ACDs among 4 individual cells for each condition. The black bar indicates the median value for each data set. For statistical test, data from 4 individual cells for each condition were pooled together and Mann-Whitney *U* test was applied. (**G**) 3D *iso*-surface reconstruction of ACDs (green) identified by using the DBSCAN algorithm for WT (left panel), CTCF depletion (middle) and RAD21 depletion (right panel) conditions. The iso-surface in grey outlines the nuclear envelope. The lower panels show 4× magnification of local regions under each condition. Scale bar, 1 μm.

Acute loss of CTCF or Cohesin triggered prominent organizational changes in ACDs (**Fig. 2B and C**) and markedly enhanced accessible chromatin clustering as the *g(r)* function increased significantly across multiple scales relative to unperturbed conditions (**Fig. 2D**). The clustering amplitude (A, equals to *g(0))*, as estimated from a fluctuation model (**Equation S11–14**), was inversely correlated with residual CTCF or RAD21 protein levels in single cells (**Fig. 2E, Fig. S7A**). More importantly, the increase in accessible chromatin clustering induced by depletion of CTCF or Cohesin was largely reversed as CTCF or RAD21 levels recovered after auxin washout (**Fig. S7, B and C**). Notably, we found that CTCF depletion preferentially increased the localization density within ACDs, whereas Cohesin removal induced the formation of much larger ACDs (**Fig. 2, F and G**). These results suggest that CTCF and Cohesin likely regulate ACD organization *via* distinct mechanisms. It is important to note that neither acute loss of CTCF nor that of RAD21 significantly affected chromatin accessibility at enhancers and promoters (**Fig. S8**). However, accessibility at CTCF binding sites was reduced upon loss of CTCF but was not impacted by Cohesin removal (**Fig. S8**). We observed similar findings by knocking down CTCF using shRNAs (**Fig. S7, D-F**). We concluded that acute loss of CTCF or Cohesin profoundly alters the 3D spatial organization of enhancers and promoters without affecting their accessibility.

### CTCF and Cohesin separate active chromosomal segments *in cis*

The enhanced accessible chromatin clustering upon CTCF or Cohesin depletion and the spatial correlation between ACDs and active chromosomal segments prompted us to examine whether CTCF and Cohesin regulate spatial segregation between active segments *in cis* (within the same chromosome) or *in trans* (from different chromosomes). To do so, we quantified physical distances between active segments in chromosomes 4 and 6 with 3D Oligopaint FISH (Beliveau et al., 2012) (**Fig. 3A**).We found that active segments *in cis* were less separated after acute loss of CTCF or Cohesin (**Fig. S9, A-C**), while distances between active segments *in trans* were largely unaffected (**Fig. S9, D and E**). These results suggested that both CTCF and Cohesin play a role in maintaining the physical separation between active segments in *cis.*

**Figure 3.**
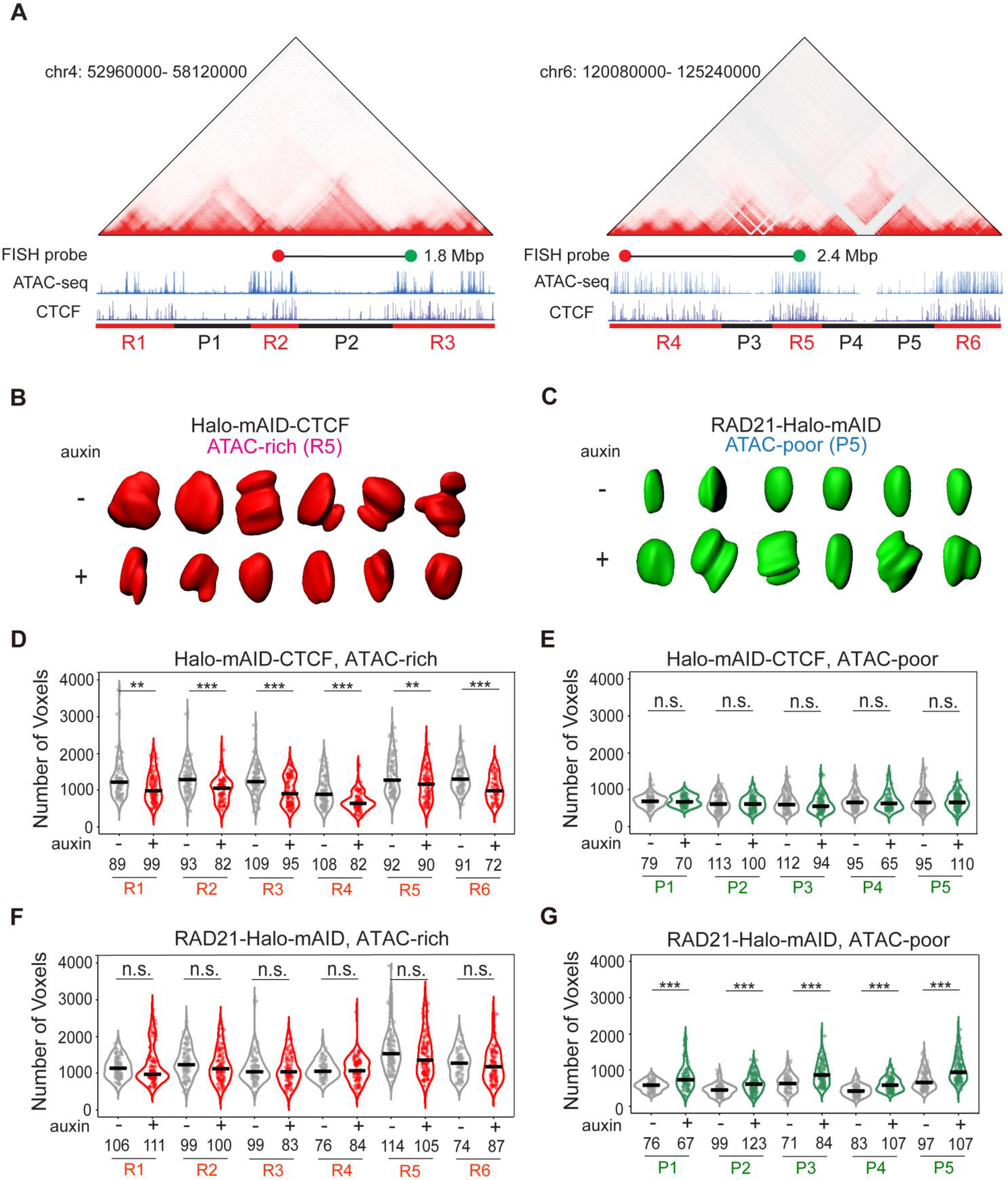
Distinct roles of CTCF and Cohesin in chromatin compaction. (**A**) Alignment of Hi-C heatmap, ATAC-seq, CTCF and RAD21 ChIP-seq tracks for two chromosomal regions harboring pluripotency genes *Klf4* (Chr4) and *Nanog* (Chr6). DNA-FISH probes corresponding to loci pairs in the adjacent ATAC-rich segments are marked as red and green dots joined by a black line. The linear genomic distance between the two loci is indicated on the side. ATAC-rich (R1-6; red) and ATAC-poor (P1-5; black) segments are underlined. (**B**) Representative iso-surfaces of an ATAC-rich segment (R5, red) before (-) and after (+) CTCF depletion. (**C**) Representative iso-surfaces of an ATAC-poor segment (P5, green) before (-) and after (+) RAD21 depletion. (**D** and **E**) Violin plot of 3D volumes (the number of voxels) of six ATAC-rich segments (R1-R6) (**D**) and five ATAC-poor segments (P1-P5) (**E**) before (grey) and after (red) CTCF depletion. The number of alleles analyzed is indicated at the bottom. The black bar indicates the median value for each data set and Mann-Whitney U test was applied for statistical test. (**F** and **G**) Violin plot of 3D volumes (the number of voxels) of six ATAC-rich segments (R1-R6) (**F**) and five ATAC-poor segments (P1-P5) (**G**) before (grey) and after (red) RAD21 depletion.

### Distinct roles of CTCF and Cohesin in chromatin compaction

Intrigued by distinct changes in accessible chromatin density and ACD size upon CTCF or Cohesin depletion, we explored whether these architectural factors differentially regulate chromatin compaction. To this end, we labeled 6 active and 5 inactive chromosomal segments with high density Oligopaint FISH probes (**Fig. 3A**). We measured the 3D volume and the sphericity score for each segment after *iso*-surface based segmentation of Airyscan images. We found that in general active segments occupy larger volumes and display lower sphericity scores relative to inactive segments (**Fig. S9, F and G**), in good agreement with STORM imaging results (Boettiger et al., 2016; Wang et al., 2016). Notably, 3D volumes of active segments were significantly reduced upon CTCF depletion, whereas inactive segments appeared unaffected (**Fig. 3, B, D and E, Fig. S9H**). In stark contrast, Cohesin depletion led to decompaction of inactive segments without causing significant changes in active segments (**Fig. 3, C, F and G, Fig. S9I**). These results revealed distinct roles of CTCF and Cohesin in the compaction of active *versus* inactive chromatin.

### Polymer modeling incorporating loop extrusion and TF-mediated weak interactions

The loop extrusion model has been used to interpret 3D genome configuration and has generated simulation data consistent with Hi-C experimental results (Fudenberg et al., 2016; Sanborn et al., 2015). To dissect physical mechanisms underlying the ACD organization, we performed loop-extrusion simulation on a 3Mb fragment consisting of one inactive segment in the middle flanked by two active segments (see **Fig. S10A** and simulation details in Supplementary Information). Remarkably, the loop extrusion model fully recapitulated the impact of CTCF depletion, including the increase in accessible chromatin clustering, the reduction in volumes of active segments, and the shortened distances observed between active segments *in cis* (**Fig. S10, A-D**). At the same time, the simulation was unable to recapitulate our experimental observations after Cohesin loss, predicting a decrease in accessible chromatin clustering, increased distances between active segments *in cis*, and decompaction of both active and inactive chromatin (**Fig. S10, A-D**).

The discrepancy between simulation and the Cohesin depletion results led us to speculate that mechanisms other than loop extrusion may be at play. Recent findings suggested that TFs and cofactors assemble in the nucleoplasm into dynamic clusters or hubs, via weak, multivalent interactions, often mediated by low-complexity domains (LCDs) (Cho et al., 2018; Chong et al., 2018; Sabari et al., 2018). To determine whether this feature contributes to ACD formation, we took advantage of the aliphatic alcohol 1,6-hexanediol (1,6-HD) that efficiently disrupts hydrophobic LCD-LCD interactions (Kato and McKnight, 2017). Low dosage 1,6-HD treatment selectively reduced accessible chromatin clustering in Cohesin-depleted cells (**Fig. 4A**), suggesting LCD mediated interactions supplied a critical function in ACD formation after Cohesin removal. At the same time, 1,6-HD treatment did not noticeably alter the strength or distribution of ATAC-seq peaks, arguing against a secondary effect associated with drug toxicity (**Fig. S11A**). Indeed, by coupling TF-mediated weak interactions inferred from 1,6-HD experiments with loop extrusion (**Fig. 4B, Fig. S11B**), the polymer simulation was able to fully recapitulate our experimental observations, including: i) enhanced clustering of accessible chromatin and shortening of distances between active segments *in cis* upon CTCF or Cohesin removal, ii) reduced clustering of accessible chromatin after 1,6-HD treatment, and iii) the volume changes in active and inactive segments after CTCF or Cohesin depletion (**Fig. S11, C and D, Fig. 4, C and D**). Results generated from the revised polymer model were insensitive to changes in TF concentrations or CTCF bead distribution (**Fig. S12, A-G**), highlighting the generality and robustness of the model. Representative chromatin configurations based on the polymer model were shown in **Fig. 4B and Movie S6**. Consistent with this integrated model (**Fig. 4E**), protein-DNA and protein-protein interactions, multivalency and optimal interaction strengths were all required to recapitulate the experimental results (**Fig. S12H**).

**Figure 4.**
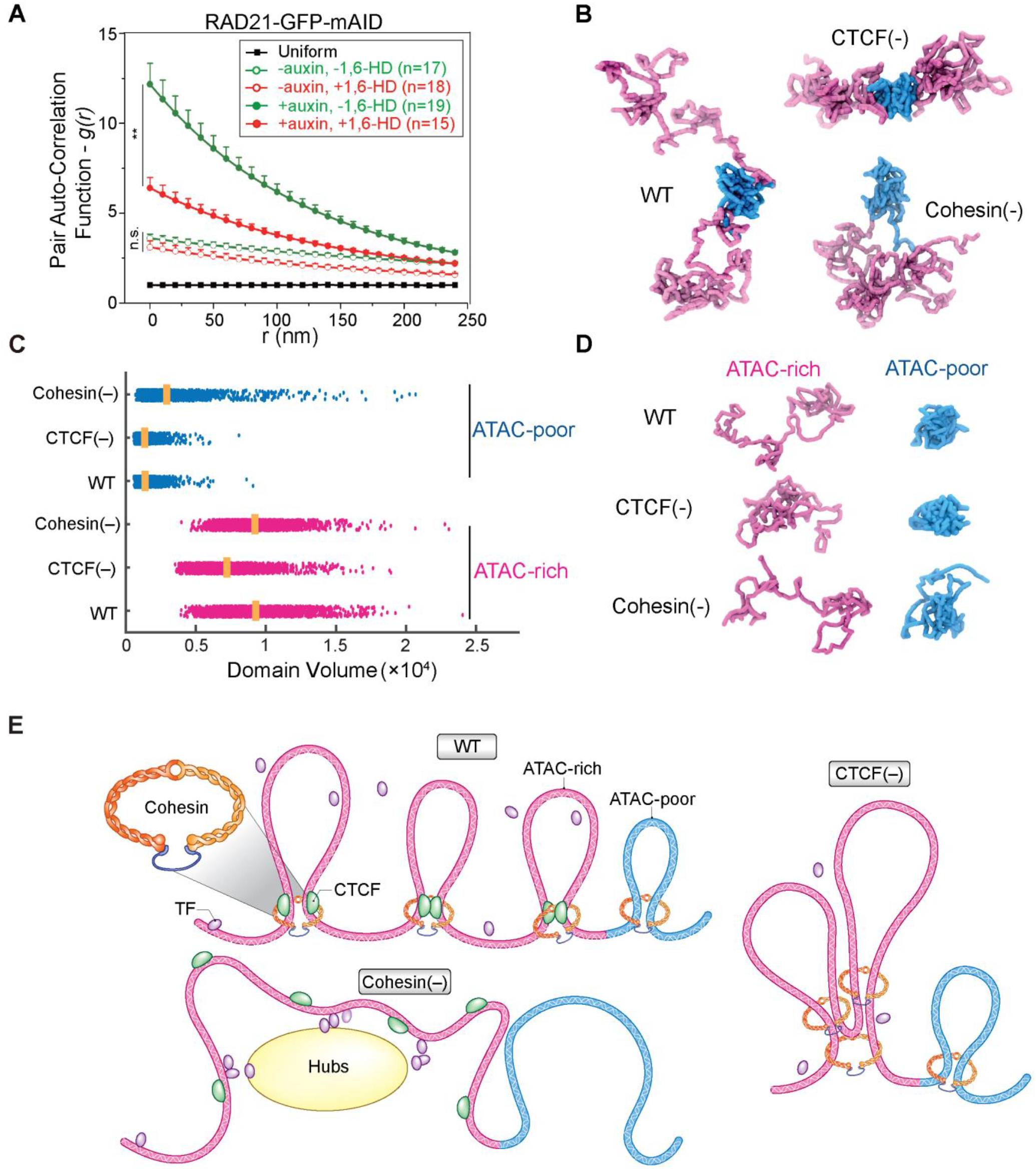
Polymer modeling with loop extrusion and TF-mediated weak interactions interprets experimental observations. (**A**) Disruption of LCD-mediated protein-protein interactions by 1,6 Hexanediol (1,6-HD) reduces accessible chromatin clustering after Cohesin depletion. *g(r)* curves were plotted for indicated conditions. Specifically, low-dose 1,6-HD (2%) treatment for 1.5 hour was applied to both control (- auxin) and Cohesin-depleted (+ auxin) cells. (**B**) Representative intra-chromatin conformation snapshots from polymer simulation incorporating both loop extrusion and TF-LCD-mediated chromatin interactions. Simulation snapshots represent WT, CTCF depletion (-) and Cohesin depletion (-). ACD flexibility is properly maintained and adjacent ACDs are well segregated in WT mESCs. CTCF depletion leads to ACD compaction and therefore the decrease of separation between ATAC-rich segments, while Cohesin depletion preferentially facilitates spatial mixing between adjacent ATAC-rich segments. (**C**) Polymer model predictions of the domain volume of ATAC-rich (red) and ATAC-poor (blue) segments under different conditions. (**D**) Representative chromatin conformation snapshots of the ATAC-rich (red) and ATAC-poor (blue) segments under normal and perturbed conditions. CTCF depletion results in ATAC-rich segment compaction, while Cohesin depletion results in ATAC-poor segment decompaction. (**E**) Illustrations of putative chromatin configurational changes under WT, Cohesin depletion or CTCF depletion conditions. ATAC-rich and ATAC-poor segments are shown in pink and cyan color, respectively. Following CTCF depletion, Cohesin continues extruding DNA fiber and progressively generates larger chromatin loops, thus effectively bringing closer two distant DNA segments and physically compacting the ATAC-rich segment. CTCF loss has minimal effect on the 3D volume of ATAC-poor segment due to lack of CTCF binding. In contrast, Cohesin depletion removes chromatin loops in both ATAC-rich and ATAC-poor segments. Although loss of chromatin loops decompacts the ATAC-poor segment, the 3D volume of the ATAC-rich segment remains largely unchanged because of the rescue effect from cooperative protein-protein and protein-DNA interactions mediated by TF hubs.

### Loss of CTCF or Cohesin alters TF dynamics in live cells

To examine whether acute loss of CTCF or Cohesin affects the organization of ACDs in live cells, we systematically mapped the 3D spatial distribution of stable SOX2 binding events in untreated, CTCF or RAD21 depleted ESCs, by using a LLSM-based imaging strategy (Legant et al., 2016; Liu et al., 2014) (**Fig. S13, A-C**). Consistent with the enhanced clustering of ATAC-PALM localizations in fixed cells (**Fig. 2D**), long-lived (>4 *sec*) SOX2 binding events also showed increased spatial clustering upon CTCF or Cohesin depletion (**Fig. 5A Movie S7**). These results are consistent with the enhanced clustering of accessible chromatin upon CTCF or Cohesin loss, because SOX2 binding sites represent a subset of accessible chromatin in the genome.

**Figure 5.**
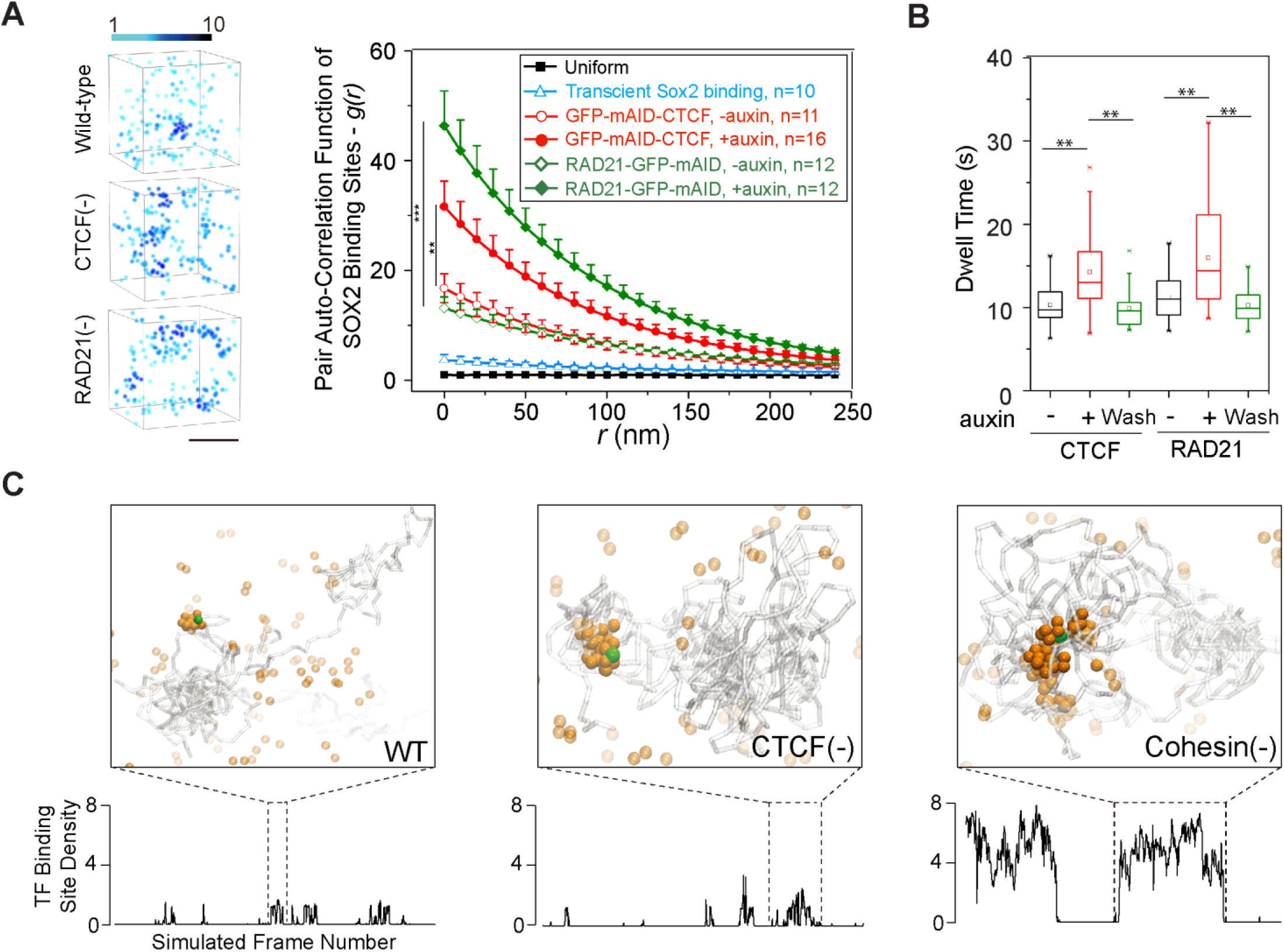
Loss of CTCF or Cohesin alters TF dynamics in live cells. (**A**) Enhanced clustering of SOX2 stable binding sites upon CTCF and RAD21 depletion. Left panel shows measured 3D spatial distribution of stable SOX2 binding sites in WT, CTCF-depleted and RAD21-depleted conditions. The local density of stable SOX2 binding sites is color coded. Scale bar, 1 μm. In the right panel, the pair auto-correlation function *g(r)* for stable SOX2 binding sites was calculated before and after CTCF or RAD21 depletion. SOX2 binding events with dwell times longer than 4s were considered as stable binding events. The error bars represent standard error of the mean. The Mann-Whitney U test was used for statistical testing. (**B**) Box plots of SOX2 dwell times before (-) and after (+) CTCF or RAD21 removal or after auxin washing-off (wash) measured by single-molecule imaging at 2 Hz. Mann-Whitney U test was performed for statistical comparison. (**C**) Simulated 3D chromatin configurations (top panel) and corresponding temporal density profiles (bottom panel) show that more TFs are “trapped” inside ACDs upon CTCF or RAD21 depletion. Simulated graphs show the moments of the tracked TF bound to chromatin. Gray lines illustrate chromatin fibers. Green sphere indicates the current position of a tracked TF. Each orange sphere represents one TF molecule that has the potential attractive interaction with binding sites on chromatin.

In order to understand the physiological function of ACDs, we next examined whether the dysorganization of ACDs affects the dynamics of TF binding and diffusion in live cells by single particle tracking (SPT). By exploiting motion blur to selectively image long-lived binding events at 2 Hz, we found that SOX2 dwelling time at specific positions became significantly elevated upon acute loss of CTCF or RAD21 and recovered to untreated levels upon auxin removal (**Fig. 5B**). Our polymer model-based MD simulations faithfully reproduced enhanced TF binding site clustering after CTCF or Cohesin loss (**Fig. S14B**). We also computed TF binding states at individual ACDs and observed two-state dynamics corresponding to the shuttling of TFs in and out of ACDs. We observed that, after CTCF or Cohesin loss, TFs formed larger clusters and resided in these clusters for a longer period of time compared to untreated cells (**Fig. 5C, Fig. S14, C-E**). These results suggest that enhanced clustering of accessible chromatin could entrap TFs for longer durations. We note that SOX2 fast diffusion in the nucleus was largely unaffected by the removal of CTCF or Cohesin, despite that target search times were slightly elevated (**Fig. S13, D-F, Fig. S14A**).

## Discussion

### ACD organization: structure and mechanism

In this study, we systematically probed the structure and organizing mechanisms of accessible chromatin in single cells, by a combination of 3D ATAC-PALM, Oligopaint FISH, genetic perturbations and polymer simulation. Consistent with previous Hi-C studies and theoretical discussions (Fudenberg et al., 2016; Nora et al., 2017; Nuebler et al., 2018; Rao et al., 2017; Sanborn et al., 2015), our data support a crucial role of CTCF and Cohesin in 3D genome organization. Gratifyingly, the loop extrusion model provided significant mechanistic insights into several of our key imaging observations. First, we found that loss of CTCF condenses active chromatin. This observation can be readily explained by the loop extrusion model as disruption of CTCF-mediated loop anchoring results in unchecked loop extrusion by Cohesin and the increased probability of extruding longer loops in accessible chromatin. Consequently, there should be a reduction in 3D volumes occupied by active chromosomal segments, consistent with the notion that loop extrusion promotes DNA compaction (Alipour and Marko, 2012; Ganji et al., 2018; Goloborodko et al., 2016). We have shown that the compaction of accessible chromatin increases localization densities in ACDs and reduces physical separation between active chromosomal segments *in cis.* Moreover, loop extrusion by Cohesin is thought to promote long-range chromatin interactions in both active and inactive chromatin domains (Fudenberg et al., 2016; Nuebler et al., 2018; Sanborn et al., 2015). When this function is blocked, inactive chromatin undergoes decondensation and expands in volume just as we observed.

Interestingly, we found that the loop extrusion model alone was insufficient to recapitulate the ACD organizational changes observed upon Cohesin depletion. Instead, the incorporation of weak TF-mediated multivalent interactions into our polymer model can faithfully reproduce our experimental observations. These results raised the intriguing possibility that genome folding requires a balanced action of loop extrusion and TF mediated protein-protein and protein-DNA/RNA interactions. Consistent with this model, a recent chromatin tracing experiment reported that globular ‘TAD-like’ domain structures persist after Cohesin depletion (Bintu et al., 2018), indicating mechanisms other than Cohesin-mediated loop extrusion may be required to establish such chromatin contacts. Because many TFs (and also cofactors, RNA Pol II *et al.*) contain intrinsically disordered LCDs that may help drive the formation of local concentrated hubs in the nucleus of live cells (Cho et al., 2018; Chong et al., 2018; Lu et al., 2018; Sabari et al., 2018), we hypothesized that dynamic, multivalent and selective interactions conferred by LCDs and orchestrated by TFs represent one contributing mechanism in mediating the organization of accessible chromatin in parallel with the CTCF- and Cohesin-driven loop extrusion. In support of this hypothesis, disruption of LCD-mediated protein hub formation by 1,6-HD severely compromised the clustering of accessible chromatin upon Cohesin removal. Moreover, a recent study found that optically activated intrinsically disordered proteins preferentially grow in regions with low densities of accessible chromatin and can mechanically restructure targeted genomic sites (Shin et al., 2018). A ‘phase separation’ mechanism has been proposed to explain heterochromatin formation (Larson et al., 2017; Strom et al., 2017) and transactivation domains of some TFs have been reported to form ‘phase separated’ liquid droplets with mediator components *in vitro* (Boija et al., 2018). The relevance of phase-separated condensates in the context of ACD organization *in vivo* will require further experimentation. To fully characterize the role of specific LCD-containing proteins in the formation of ACDs, it will be important to identify a live-cell ACD marker and to perform imaging experiments to study the spatiotemporal dynamics of ACDs and their functional relationship to TF hubs.

### Functional impact of ACDs

The gene-rich *A* compartment was detected by Hi-C 10 years ago (Lieberman-Aiden et al., 2009). However, its spatial distribution and organizational pattern have not been directly visualized in single cells. Because the *A* compartment can be computationally deduced from sequencing-based chromatin accessibility data (Buenrostro et al., 2015; Fortin and Hansen, 2015), we speculate that ACDs, uncovered for the first time by 3D ATAC-PALM imaging, may represent the physical organization of the *A* compartment (or a portion of it). The preferential spatial mixing of ACDs upon Cohesin removal is consistent with the enhanced compartmentalization and increased super-enhancer contacts after depleting Cohesin or its loading factor (Nuebler et al., 2018; Rao et al., 2017; Schwarzer et al., 2017). Interestingly, CTCF loss exhibited a weaker effect in spatial mixing of ACDs but a more pronounced impact in ACD compaction. This observation agrees with the proposed role of CTCF in insulating the TAD and loop domains based on Hi-C studies (Kubo et al., 2017; Nora et al., 2012; Wutz et al., 2017). It is important to note that since CTCF-mediated chromatin folding is dose-dependent on its nuclear concentration (Nora et al., 2017), our near-complete CTCF depletion based on the N-terminal tagging strategy could potentially induce more pronounced changes in genome organization than what was reported for C-terminal AID mediated degradation systems (*33, 42*).

The clustering of ATAC-PALM localizations highlights the prevalent existence of spatial proximity among *cis-* regulatory elements. This observation is reminiscent of the extensive promoter-promoter, enhancer-enhancer and enhancer-promoter interactions captured by ensemble chromatin interaction experiments (Kieffer-Kwon et al., 2013; Li et al., 2012). We imagine that the local high-density clustering of regulatory DNA elements would increase the local concentration of TF and coactivators, promote cooperative protein-chromatin and protein-protein interactions, facilitate diffusion to proximal regulatory DNA elements, ensnare TFs binding and allow multiple rounds of local target search cycles. Consistent with this model, live-cell single-molecule imaging and molecular simulation showed that enhanced accessible chromatin clustering upon CTCF or Cohesin loss is accompanied by a more clustered distribution of TF stable binding events in the nucleus and increased TF dwell times in individual ACDs. The architecture of clustered *cis-* regulatory elements may thus serve as a topological basis for mediating long-range enhancer-promoter communications. It is more likely that the transient spatial proximity among TFs and cofactors within locally concentrated hubs, rather than the stable tethering among them, orchestrates gene activation. In the future, high-throughput, sequence-specific Oligopaint FISH-based barcoding strategies in combination with 3D ATAC-PALM may help delineate the spatial relationship between ACDs, compartments, TADs and loops (Bintu et al., 2018; Chen et al., 2015; Shah et al., 2018; Wang et al., 2016). Optogenetic tools with high spatiotemporal resolution will deepen our mechanistic understanding of ACD formation (Shin et al., 2018). These strategies, when coupled with MS2 or smFISH based gene activity measurements, might address the fundamental yet puzzling question of how local genome organization and transient protein hubs contribute to transcriptional control.

## MATERIALS AND METHODS

### Tn5 Expression and Purification

The plasmid pTXB1-Tn5 (Addgene #60240) was used to express the hyperactive Tn5 transposase fused with the Mxe Intein and Chitin-binding domain (Picelli et al., 2014). Briefly, the pTXB1-Tn5 was transformed into the T7 Express lysY/Iq Competent E. coli (NEB, cat# C3013I). 1 litter culture of transformed cells were grown in TB buffer with antibiotics at 37°C to A600 ~0.6 and induced with 0.25 mM IPTG. The cells were incubated at 23°C for 4 h, harvested, and washed with 50 mL 1 × PBS with complete^™^ Protease Inhibitor Cocktail (Roche). The cell pellet was snap frozen in liquid nitrogen, stored at −80°C overnight, resuspended in 160 mL HEGX (20 mM HEPES-KOH at pH 7.2, 0.8 M NaCl, 1 mM EDTA, 10% glycerol, 0.2% Triton X-100) with Protease Inhibitor and sonicated for 10 cycles of 30s ON/60s OFF at 30 W output on a Model 100 Sonic Dismembrator (Fisher Scientific). The lysate was centrifuged in a Beckman JA25.50 rotor at 16,000 rpm for 30 min at 4°C and 8.4 mL 5% neutralized PEI (Sigma P3143) was added dropwise into the supernatant with gentle agitation for 15 minutes at 4°C followed by another centrifuge at 13,000 rpm for 10 min at 4°C. Meantime 20 mL Chitin Resin (NEB cat#S6651L) was equilibrated in a column with 200mL of HEGX buffer supplemented with protease inhibitor and loaded with the supernatant from the PEI-centrifuge step by slow gravity flow (<0.3mL/min). The column was washed with 400 mL HEGX overnight at a flow rate ~0.6 mL/min. To cleave the Tn5 from the Intein, 48 mL HEGX buffer + 100 mM DTT supplemented with protease inhibitor was loaded to the top of the column bed and the column was left closed for 48 h at 4° C. Cleaved Tn5 was eluted in ~1 mL fractions and Tn5 elution efficiency was monitored by the Pierce^™^ Coomassie (Bradford) Protein Assay Kit (Thermo Fisher Scientific cat# 23200). Fractions with the strongest absorbance were pooled and dialyzed versus two changes of 2 L of 2X Tn5 dialysis buffer (100 mM HEPES-KOH at pH 7.2, 0.2 M NaCl, 0.2 mM EDTA, 2 mM DTT, 0.2% Triton X-100, 20% glycerol) at 4° C. After dialysis, the protein concentration was measured using a Nanodrop spectrophotometer (Thermo Scientific) and confirmed by running an SDS-PAGE gel using BSA as a standard. Tn5 protein was further concentrated to 20 mg/mL using the Amicon Ultra-15 Centrifugal Filter Unit and we added 50% glycerol to the final concentrate and prepared aliquots in screw-top microcentrifuge tubes, flash freeze in liquid nitrogen, and store at −80° C. The Tn5 activity was tested by the *in vitro* tagmentation assay (Picelli et al., 2014) and further verified by performing genome wide ATAC-seq experiment.

### PA-JF_549_ dye-oligo conjugation and purification

The mosaic ends (ME) adaptors for Tn5 transposase were synthesized by Integrated DNA Technologies (IDT) with attachment of a 5’ end primary amino group by a six-carbon spacer arm (C6). The oligonucleotide sequences and their modifications are:

Tn5ME-A: 5’-amino-C6 TCGTCGGCAGCGTCAGATGTGTATAAGAGACAG3’

Tn5ME-B: 5’-amino-C6-GTCTCGTGGGCTCGGAGATGTGTATAAGAGACAG-3’

Tn5MErev: 5’-[phos]CTGTCTCTTATACACATCT-3’

The photoactivatable Janelia Fluor 549 coupled to the N-hydroxysuccinimide (NHS) ester (NHS-PA-JF_549_) was synthesized by the Luke Lavis’s group at Janelia Research Campus. The NHS-PA-JF_549_ was dissolved in anhydrous dimethylsulfoxide (DMSO) immediately before conjugation. The Tn5ME-A and Tn5ME-B oligos were first dissolved in deionized water and then extracted three times with chloroform. The upper aqueous solution was carefully extracted and precipitated by 3M sodium acetate (pH 5.2) and ethanol. The pellet was washed with 70% ethanol, dried and dissolved in ultrapure water. The purified oligos were reacted with excessive NHS-PA-JF_549_ in anhydrous DMSO (mass ratio 1:2.5) in the labeling buffer (0.1 M sodium tetraborate, pH 8.5). The reaction was incubated at room temperature for overnight (>12 hours) with constant stirring and protection from light. The conjugation reaction was further precipitated by 3M sodium acetate, pH 5.2 and ethanol. The purified pellet was dissolved in 0.1 M TEAA (triethylammonium acetate) (Thermo Fisher Scientific cat# 400613).

Purification of oligo-dye conjugates were performed on Agilent 1200 Analytical HPLC system equipped with an autosampler, diode array detector and fraction collector. The column used was Eclipse XDB-C18 column (4.6 x 150 mm 5 μm, Agilent) and eluted with a linear gradient of 0-100% MeCN/H2O with constant 10 mM TEAA additive; 30 min run; 1ml/min flow, detection at 260 nm. Sample fractions were pooled and lyophilized to obtain the product as white solid.

### *In vitro* Tn5 transposome assembly, validation and preparation for PALM imaging

The HPLC purified Tn5ME-A-PA-JF_549_, Tn5ME-B-PA-JF_549_, Tn5MErev oligos were resuspended in water to 100 μM each. Tn5ME-A-PA-JF_549_ or Tn5ME-B-PA-JF_549_ were mixed with Tn5MERev in each molar ratio in 1X annealing buffer (10mM Tris HCl pH8.0, 50mM NaCl, 1mM EDTA) and were denatured on a benchtop thermocycler at 95 °C for 5 min and then slowly cooled down to 25°C at the rate of −1°C /min. We assembled the Tn5 transposome with PA-JF_549_ according to previously published protocol (Chen et al., 2016) by combining 0.25 vol Tn5MErev/Tn5ME-A-PA-JF_549_ + Tn5MErev/Tn5ME-B-PA-JF_549_ (50 μM each), 0.4 vol glycerol (100% solution), 0.12 vol 2× dialysis buffer (100 mM HEPES–KOH at pH 7.2, 0.2 M NaCl, 0.2 mM EDTA, 2 mM DTT, 0.2% Triton X-100, 20% glycerol), 0.1 vol Tn5 (50 μM), 0.13 vol H2O, followed by gentle nutation at room temperature for 1 hour. The activity of in-house prepared Tn5 transposome was validated by genome-wide ATAC-seq as described previously (Chen et al., 2016) following an reverse crosslinking step.

We prepared cells for 3D ATAC-PALM experiments as reported previously (Chen et al., 2016). Briefly, cells (mouse ESC or MEF) were plated onto #1 thickness 5mm coverslips (Warner Instruments, cat#64-0700) at around 70-80% confluency with proper coating one day before experiment. Cells were fixed with 1% paraformaldehyde (Electron Microscopy Sciences, Cat# 15710) for 10 min at room temperature. After fixation, cells were washed three times with 1 X PBS for 5 minutes and then permeabilized with ATAC lysis buffer (10 mM Tris–Cl, pH 7.4, 10 mM NaCl, 3 mM MgCl2, 0.1% Igepal CA-630) for 10 min at room temperature. After permeabilization, the slides were washed twice in 1XPBS and put inside a humidity chamber box at 37 °C. The transposase mixture solution (1× Tagmentation buffer-10mM Tris-HCl, pH 7.6, 5mM MgCl2, 10% dimethylformamide, 100 nM Tn5-PA-JF_549_) was added to the cells and incubated for 30 min at 37 °C inside the humidity chamber. After the transposase reaction, slides were washed three times with 1 X PBS containing 0.01% SDS and 50 mM EDTA for 15 min at 55 °C before mounted onto the Lattice light-sheet microscope (LLSM) slot for 3D ATAC-PALM imaging.

### ATAC-seq libraries preparation and genomic analysis

ATAC-seq libraries were made according to the published protocols (Buenrostro et al., 2013; Corces et al., 2017) using the Nextera DNA Library Preparation Kit (illumine, cat# FC-121-1030) or home-made Tn5 transposon as described in (Picelli et al., 2014). We also supplemented 0.05ng drosophila genomic DNA purified from S2 cells right before adding the tagmentation mixture to improve comparison between samples. Multiplexed ATAC-seq libraries (barcodes were adapted from (Buenrostro et al., 2013)) were sequenced on Illumina HiSeq 2500 or NextSeq 500/550 high and/or mid throughput 150 cycles at Janelia quantitative Genomics Core, with a run configuration of 75 bp paired-end sequencing, either as single indexed or dual indexed runs. All samples were quantitated on Roche 480 lightcycler using FAST qPCR program and normalized and pooled at 2nM. Libraries were loaded at varied final concentrations across HiSeq (10pM) and NextSeq (1.5pM to 1.9pM). Illumina’s Bcl2tofastq2 v2.17 was used to convert BcL files to fastq files and to demultiplex the samples.

To analyze ATAC-seq libraries, pair-end reads were first adapter removed by Cutadapt and mapped to mm10 genome build using Bowtie2 with the following parameters: --no-discordant --no-mixed --phred33 −X2000 −threads32. Reads mapped to mitochondria/uncharacterized chromosomes (chrM/chrUn/random) and PCR duplicates were removed by samtools. The pair-end reads from the drosophila genomic DNA was mapped to dm6 genome build. To compare the coverage among ATAC-seq libraries, sequencing reads were normalized to 1×sequence depth defined by total number of mapped reads × fragment length / effective genome size (2,150,570,000) or by the spike-in drosophila genomic DNA reads. ATAC-seq peaks were called using the MACS2 callpeak function using the −f BAMPE parameter. Fragment length distribution, TSS enrichment, scatter plots were analyzed as previously described (Chen et al., 2016). Raw sequencing data were deposited to NCBI GEO with the accession number of GSE126112.

CTCF and RAD21 ChIP-seq data sets from wild type ESCs were obtai ed from (Hansen et al., 2017) under GSE number GSE90994. The Transcription start site (TSS) annotation was downloaded from the UCSC Table Browser mouse mm10 build (GRCm38/mm10, Dec.2011) NCBI RefSeq genes and the promoters were defined as −400bp to +100bp relative to the TSS. ESC enhancer coordinates were retrieved from the H3K27ac ChIP-seq peak regions as described previously (Whyte et al., 2013).

### Cell culture and generation of auxin-inducible ESC lines

JM8.N4 mouse ESCs from the C57BL/6N strain and their genome edited derivatives were cultured on 0.1% gelatin coated plates without feeders at 37°C and 5% CO_2_. The ESC medium was composed of the knockout DMEM(1X) optimized for ESCs (Thermo Fisher Scientific 10829-018), 15% ESC qualified Fetal Bovine Serum (ATCC SCRR-30-2020), 1 mM GlutaMAX (Thermo Fisher Scientific 35050-061), 0.1 mM MEM nonessential amino acids (Thermo Fisher Scientific 11140-50), 0.1 mM 2-mercaptoethanol (Thermo Fisher Scientific 21985-023),1000 units of Leukemia inhibitory factor (LIF) (Millipore) and Antibiotic-Antimycotic (Thermo Fisher Scientific 15240-062). The JM8.N4 cells were approved by the NIH 4D Nucleome project as a Tier2 cell line. Mouse embryonic fibroblast (MEF) cells were cultured in DMEM (Thermo Fisher Scientific) supplemented with 10% Fetal Bovine Serum and 1× Antibiotic-Antimycotic.

To implement the auxin-inducible degron system (AID), the pLenti-EF1-osTir1-9myc-P2A-Bsd lentivirus was used to infect the low passage JM8.N4 wild type ESCs and selected with 10μg/mL Blasticidin (Thermo Fisher Scientific R21001). Blasticidin resistant clones were manually picked and tested for osTir1 expression by Western Blot using an anti-Myc antibody (Santa Cruze, 9E10/sc-40). Cell cycle kinetics and ATAC-Seq analysis of Tir1 ESCs showed no discernable difference to parental JM8.N4 ESCs.

To generate auxin inducible degradation for CTCF and Cohesin, 1.5 μg/μl wild type SpCas9_sgRNA_PGK-Venus construct and 3 μg/μl of CTCF or RAD21 donor constructs were nucleofected into ~3X10^6 Tir1 ESCs using the Amaxa^™^ 4D-Nucleofector and the P3 Primary Cell 4D-Nucleofector^™^ X Kit following the manufacture’s protocol. ~24 hours following nucleofection, Venus positive cells were FACS sorted as a pool and grown for another ~3-5 days. Cells were then stained with 50nM Janelia Fluor^®^ 549 HaloTag ligand (JF_549_) for 30min, washed 3X with ESC medium for 15min, and subjected to another round of FACS sorting. JF_549_ positive cells were plated sparsely at 10cm tissue culture plates and grown for another 5-7 days. Single colonies were picked for genotyping by designing PCR primers outside of the homology arms. Bi-allelic knock-in clones were verified by Sanger Sequencing and expanded for downstream analysis. Genotyping primers for *Ctcf* knock-in are: 5’-CCGCCCAGTCATTTCACCTACA-3’, 5’- GGCCGTTCTGGAGTGGTTTACG- 3’; for *Rad21* knock-in are: 5’-CAGGTATGCCAGCACAGTCCACA-3’, 5’- CCAGGAATACAAACCCAACCCAAA-3’.

The GFP version of CTCF/RAD21-AID ESCs were similarly generated except that ESCs were sorted for GFP fluorescence ~5 days after initial sorting for Venus signal. Biallelic knock-in ESC clones were verified by PCR genotyping, Sanger Sequencing, and WB. To generate the stable SOX2-HaloTag expressing ESCs, 2μg PiggyBac_EF1_HaloTag_SOX2 was co-transfected with 1 μg PiggyBac super-transposase and selected with 500 μg/mL G418 (Thermo Fisher Scientific cat# 10131035) for ~7 days.

We tested a wide range of auxin concentration for target degradation and found that 50-500μM IAA induced robust CTCF or RAD21 degradation (data not shown). Unless indicated, we used 100μM final concentration of IAA throughout our experiments.

### Chemicals and plasmids

The plant auxin analog indole-3-acetic acid (IAA) (Sigma, Cat# I3750-5G-A) was dissolved in Ethanol at a stock concentration 500mM and aliquoted to store at −20°C. 1,6 Hexanediol (Sigma 240117-50G) were dissolved in ESC medium at final concentration 2% and treated 1.5 hours.

The pLenti-EF1-osTir1-9Myc-P2A-Bsd was constructed by PCR amplifying the Oryza Sativa Tir1 (osTir1) cDNA from the pBabe Puro osTIR1-9Myc (Addgene #80074) and inserted into the AgeI/BamHI site of the lentiCas9-Blast construct (Addgene #52962).

The donor plasmids for CTCF and RAD21 were modified from previous HaloTag-CTCF and RAD21-HaloTag donor constructs as described from (Hansen et al., 2017) by in-frame insertion of the 71-104aa of full length auxin-inducible degron (miniAID) (Morawska and Ulrich, 2013)(Nora et al., 2017). The CRISPR/Cas9 and sgRNA constructs used for targeted CTCF and RAD21 knock-in were used as previously described (Hansen et al., 2017). The corresponding GFP version of CTCF and RAD21 donor constructs were generated by replacing the HaloTag with mNeonGreen and eGFP, respectively, by Gibson assembly. Maps of plasmids used in this study are available upon request.

The PiggyBac_EF1_HaloTag_SOX2_IRES_neo and the PiggyBac super-transposase were used as previously described (Chen et al., 2014b). The PiggyBac_EF1_mNeonGreen_HP1α construct was made by inserting the mNeonGreen and HP1α gBlock into the PiggyBac_EF1_HaloTag_SOX2 backbone by Gibson Assembly (NEB, E5520S). The PiggyBac_EF1_H2B-GFP plasmid was made as previously described (Teves et al., 2016).

### RNAi

Lenti-viral vectors expressing shRNA against mouse CTCF and preparation of lentiviral particles were described in our previous report (Liu et al., 2011). Two different lentiviral vectors expressing CTCF shRNAs and an empty vector control were used to infect CTCF-HaloTag ESCs (Hansen et al., 2017) in the presence of 5ng/mL Polybrene (Millipore, cat# TR-1003-G). One day after infection, cells were selected with 1 μg/mL Puromycin (Thermo Fisher Scientific, cat# A1113803) for additional 48 hours before analysis with immunofluorescence, ATAC-seq and 3D ATAC-PALM microscopy. The endogenously HaloTag labeled CTCF was labeled with 100nM JF646 HaloTag ligand (Grimm et al., 2015) for 30min to monitor the degree of CTCF depletion by shRNA.

### Western Blots

ESCs were lysed in 1XSDS sampling buffer (200mM Tris HCl pH7, 10% glycerol, 2% SDS, 4% beta-mercaptoethanol, 400mM DTT, 0.4% bromophenol blue) preheated at 95°C. Lysates were further sonicated and denatured at 95°C for 5min. Proteins from each sample was resolved by SDS-PAGE using Mini-PROTEAN^®^ TGX^™^ Precast Gels (Biorad). Primary antibodies used include: CTCF (Millipore, cat#07729), RAD21 (abcam, Cat#154769), OCT4 (Santa Cruz, sc-5279), SOX2 (Millipore, Cat# AB5603). We used HRP conjugated secondary antibodies (Pierce) at a dilution of 1:3000. Western blot was exposed to Western Lightning Plus-ECL (PerkinElmer) and imaged in a ChemiDoc MP (Bio-Rad) detection system.

### Confocal Imaging and analysis

Immunofluorescence and HaloTag JF_549_ imaging for CTCF or RAD21 depletion was performed under the Nikon A1-R confocal system using the Galvo scanning mechanism. The antibodies used for immunofluorescence are SOX2 (Fisher R&D Systems, Cat# AF2018), OCT4 (Santa Cruz, Cat# C-10). We used the default excitation lines: 405/488/561/640 nm lasers and the 4 channel detector--Ch1 450/50 DAPI (multi-alkali), Ch2 525/50 GFP (GaAsP), Ch3 600/50 RFP (GaAsP), Ch4 685/70 Cy5 (multi-alkali). Multiple z-stack images (300nm) were acquired and the fluorescence intensity was obtained from the maximum intensity projection under ImageJ/Fiji.

### Colony Formation Assay and Cell proliferation analysis

Colony forming assays were performed by plating 600 cells per well on 12-well plates (150/cm^2^) coated with gelatin. Plates were fixed after up to 7 days of IAA treatment upon initial plating and stained for alkaline phosphatase (AP) activity (Sigma, cat. 86R-1KT) according to the manufacturer’s protocol.

CTCF or RAD21 depleted ESCs were assayed for DNA replication by the Click-iT^®^ EdU Alexa Fluor^®^ 488 Flow Cytometry Assay Kit (Thermo Fisher Scientific, cat#C-10420) according to the manufacture’s protocols. Prior to Flow Cytometry analysis, DNA was stained with 1μg/mL FxCycle^™^ Violet stain (Thermo Fisher Scientific, cat#F10347) at room temperature for 30min. DNA synthesis was measured by incorporating 5-ethynyl-2’-deoxyuridine (EdU) coupled with an alkyne group, followed by the click reaction with Alexa Fluor 488 dye coupled with the picolyl azide group. Flow Cytometry was performed on the CytoFLEX S system from Beckman and analyzed by FlowJo.

### 3D ATAC-PALM image acquisition and processing

The 3D ATAC-PALM data were acquired by the Lattice light-sheet microscopy (Chen et al., 2014a) at room temperature. The light sheet was generated from the interference of highly parallel beams in a square lattice and dithered to create a uniform excitation sheet. The inner and outer numerical apertures of the excitation sheet were set to be 0.44 and 0.55, respectively. A Variable-Flow Peristaltic Pump (Thermo Fisher Scientific) was used to connect a 2L reservoir with the imaging chamber with 1×PBS circulating through at a constant flow rate. Labelled cells seeded on 5mm coverslips (Warner Instruments) were placed into the imaging chamber and each image volume includes ~100-200 image frames, depending on the depth of field of view. Specifically, spontaneously activated PA-JF_549_ dye were initially pushed into the fluorescent dark state through repeated photo-bleaching by scanning the whole image view with a 2W 560 nm laser (MPB Communications Inc., Canada). Then, the samples were imaged by iteratively photo-activating each plane with very low intensity 405 nm light (<0.05 mW power at the rear aperture of the excitation objective and 6W/cm^2^ power at the sample) for 8 ms and by exciting each plane with a 2W 560 nm laser at its full power (26 mW power at the rear aperture of the excitation objective and 3466 W/cm^2^ power at the sample) for 20 ms exposure time. The specimen was illuminated when laser light went through a custom 0.65 NA excitation objective (Special Optics, Wharton, NJ) and the fluorescence generated within the specimen was collected by a detection objective (CFI Apo LWD 25×W, 1.1 NA, Nikon), filtered through a 440/521/607/700 nm BrightLine quad-band bandpass filter (Semrock) and N-BK7 Mounted Plano-Convex Round cylindrical lens (f = 1000 mm, Ø 1”, Thorlabs), and eventually recorded by an ORCA-Flash 4.0 sCMOS camera (Hamamatsu). The cells were imaged under sample scanning mode and the dithered light sheet at 500 nm step size, thereby capturing a volume of ~25 μm × 51 μm × (27~54) μm, considering 32.8° angle between the excitation direction and the stage moving plane.

To precisely analyze the 3D ATAC-PALM data, we embedded nano-gold fiducials within the coverslips for drift correction as previously described (Legant et al., 2016). ATAC-PALM Images were taken to construct a 3D volume when the sample was moving along the “s” axis. Individual volumes per acquisition were automatically stored as Tiff stacks, which were then analyzed by in-house scripts written in Matlab. The cylindrical lens introduced astigmatism in the detection path and recorded each isolated single molecule with its ellipticity, thereby encoding the 3D position of each molecule relative to the microscope focal plane. All processing was performed by converting all dimensions to units of xy pixels, which were 100 nm × 100 nm after transformation due to the magnification of the detection objective and tube lens. We estimated the localization precision by calculating the standard deviation of all the localizations coordinates (x, y and z) after the nano-gold fiducial correction. The localization precision is 26±3 nm and 53±5 nm for xy and z, respectively.

Raw 3D ATAC-PALM images were processed following previous published procedures (Chen et al., 2014a; Legant et al., 2016). Specifically, a background image with gray values taken without illumination was subtracted from each frame of the current raw image. Photo-activated images were then filtered by subtracting Gaussian filtered images at two different standards:

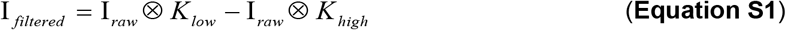

Where ⊗ is the convolution operator, and *K_low_* and *K_high_* represent 5×5 pixel two-dimensional Gaussian function with standard deviation σ=2 (low) and 1 (high):

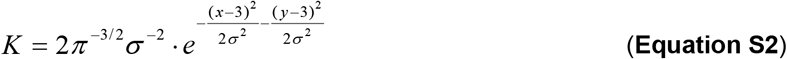

The local maxima were then determined on these filtered images and the coordinates were estimated and used as the true position of the isolated single molecule.

In order to precisely determine the z position of each isolated localization, cylindrical lens were used and the corresponding astigmatic PSF function could be formulated as a Gaussian function with separate x and y deviations. Importantly, these deviations encoded the relative z-offset of the signal with respect to the focal plane.

Where

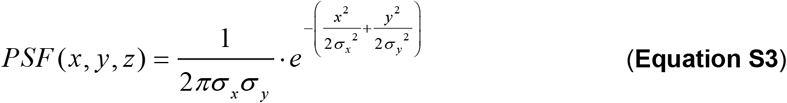

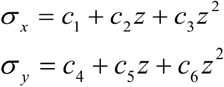

Here *σ_x_* and *σ_y_* were polynomial function of z. The constants *c_1_~c_6_* can be fitted by scanning bright and regular beads through the focal plane using piezoelectric stage.

To rigorously compare ATAC-PALM localizations within the nucleus, image stack for H2B-GFP stably expressed in mouse ESCs was captured before the ATAC-PALM experiment under the same Lattice light-sheet Microscopy imaging condition. The image stack of the GFP channel was de-skewed and de-convoluted using the same algorithm to identify a nuclear mask to segment ATAC-PALM signal. The percentage of localizations falling into the same volume in or out of the mask were counted and plotted in Figure S2B. We found that the majority (>95%) of localizations fall into the nuclear H2B mask, suggesting that the manual segmentation based on ATAC-PALM signal intensity is sufficient to distinguish it from out-of-nucleus signal (e.g. mitochondria) or non-specific background.

To derive the ATAC-PALM intensity map in order to compare with HP1-GFP labeled heterochromatin spatial distribution, ATAC-PALM localizations were binned within a cubic of 100 nm with a 3D Gaussian filter and a convolution kernel of 3×3×3.

### 3D Imaging of SOX2 enhancer clusters in live cells

ESCs stably expressing HaloTag-SOX2 in the CTCF/RAD21-GFP-AID genetic background were plated onto #1 5mm round coverslips (Warner Instruments, 64-0700, CS-5R) pre-coated with Corning Cell-Tak matrix (Corning, 354240). Cells were stained with HaloTag ligand JF_549_ at a final concentration of 10 nM for 15 mins, washed in PBS twice before being mounted onto the sample holder. The imaging medium was prepared by supplementing FluoroBrite DMEM Medium (Thermo Fisher Scientific) with 15% ES Cell Qualified Bovine Serum, 1 mM Glutamax, 0.1 mM nonessential amino acids, 1 mM sodium pyruvate, 0.1 mM 2-mercaptoethanol, and 1000 units of LIF. Before experiments, the LLSM was calibrated with imaging medium at 37 °C overnight. During experiments, the imaging chamber was filled with 8 mL imaging medium containing 40pM HaloTag ligand JF_549_. All live cell samples were imaged by iteratively exciting each plane with a 560 nm laser (~10 mW power at the rear aperture of the excitation objective) at 50 ms exposure time. The sample cells were imaged around 200 frames per sample volume by sample scanning mode with the dithered light sheet of 500 nm step size, thereby capturing a volume of ~25 μm × 51 μm × 54 μm.

3D localization (x, y, z) of detected single molecules was further analyzed as previously described (Liu et al., 2014). The PSF model can be described by the following equation:

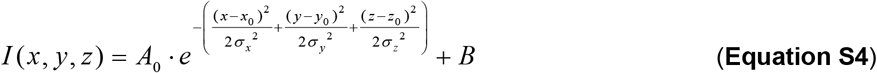

Where *A_0_* is the signal amplitude and *B* is the background signal value.

Image registration and drift correction were similarly performed for the ATAC-PALM methods. The centroid displacement of total localization events from every 40 time points (160 s) was calculated and the resulting transformation matrix over time was applied to the data accordingly. Significant drifted datasets were eliminated in the following tracking analysis.

### 3D pair auto-correlation function and *Ripley’s* function

As described previously (Liu et al., 2014), the pair correlation function *g*_0_ (*r*) or radial distribution function measures the probability *P* of finding a localization of accessible chromatin in a volume element *dV* at a separation *r* from another accessible chromatin site:

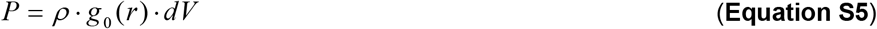

Where ρ represents the mean density of accessible chromatin in the nucleus. 3D *g*_0_ (*r*) was computed by

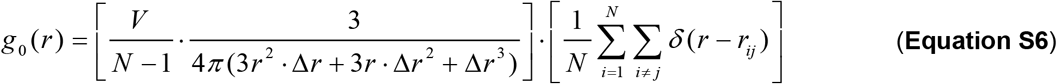

N is the total number of localizations and (N-1)/V is the average localization density within the 3D nuclear volume *V*. △*r* = 50 nm is the binning width used in the analysis. The item 4*π*(3*r*^2^ · Δ*r* + 3*r* · Δ*r*^2^ + Δ*r*^3^)/3 represents the shell-shape volume between the search radius from *r* to *r* + Δ*r*. The Dirac Delta function is defined by

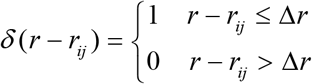

Where *r_ij_* represents the pair-wise Euclidean distance between localization point *i* and *j.* To eliminate the boundary effect in calculation of *g*_0_(*r*)from a finite 3D volume, we generated *g_r_* (*r*) from uniform distributions with the same localization density in the same volume as real data using the Matlab convex hull function. Accordingly, the final normalized 3D pair auto-correlation function *G*(*r*) was calculated by:

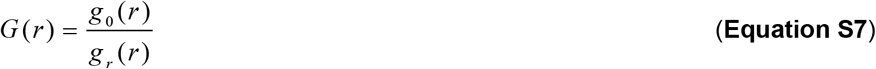

Ripley’s K function is a variation of pair correlation function to study point distribution patterns and is defined by the equation:

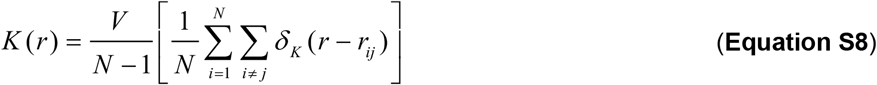

where its Delta function given by

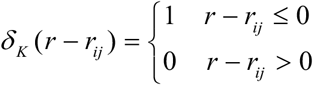

The *K*(*r*) for a random Poisson distribution is *πr*^2^. Therefore, its derivative Ripley’s L function is given by

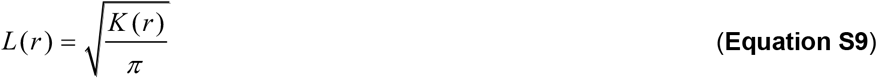

The deviation of *L*(*r*) from expected random Poisson distribution can be used to indicate point clustering or dispersion. The further normalized Ripley’s H function

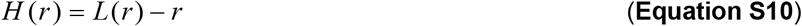

Can be used to estimate the point clustering and domain size (Kiskowski et al., 2009).

Whereas *G*(*r*) can be used as *bona fide* criteria for estimating the clustering effect of a group of spatial localizations, it has to be adjusted regarding the inevitable localization uncertainty of the PSF. If the distribution is purely random the revised pair auto-correlation *G_R_* (*r*) considering PSF uncertainty is given by

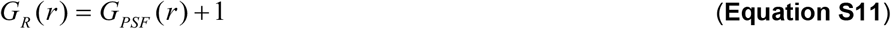

with

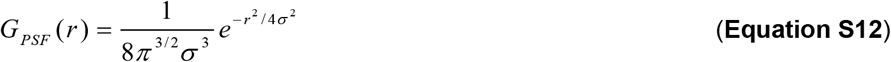

where *ρ* represents the average surface density of molecules, *G_PSF_*(*r*) denotes the correlation of uncertainty in the PSF and *σ* for standard deviation. The spatial autocorrelation of accessible chromatin can be approximated by an exponential function:

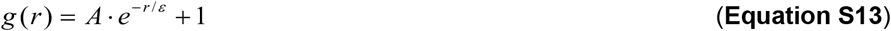

where *ε* and A denotes the correlation length and amplitude of the accessible chromatin cluster, respectively. The total correlation can be described by

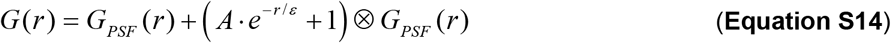

The raw data were fitted by using Equation S14 to derive *g*(*r*) and parameters *A* and ε. The approximated function *g*(*r*) was used throughout the paper for pair auto-correlation analysis. Curve fitting was performed using the trust-region method implemented in the Curve Fitting Matlab toolbox.

### 2D pair cross-correlation function

To calculate the spatial cross-correlation between the two color ATAC localization signal and the heterochromatin HP1α signal, we first converted the localization densities into intensity map via 2D Gaussian blurring. The pair cross-correlation function 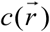 was then calculated by the fast Fourier transform (FFT) method described previously (Liu et al., 2014).

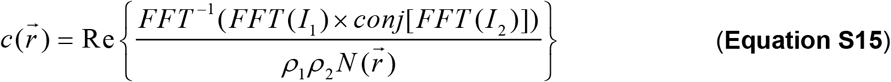

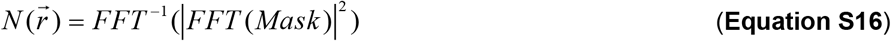

The 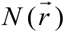 is the auto-correlation of a mask matrix that has the value of 1 inside the nucleus used for normalization. *conj*[] refers to complex conjugate, ρ_1_ and ρ_2_ are the average surface densities (number of localizations divided by the surface area) of images I_1_ and I_2_, and Re{} indicates the real part. The fast Fourier transform and its inverse (*FFT* and *FFT*^−1^) were computed by fft2() and ifft2() functions in Matlab, respectively. Cross correlation functions were calculated first by converting the Cartesian coordinates to polar coordinates by Matlab cart2pol() function, binning by radius and by averaging within the assigned bins.

### DBSCAN analysis

The density-based clustering algorithm DBSCAN (Density-Based Spatial Clustering of Applications with Noise) was adopted to map and visualize individual local ACDs (core DBSCAN Matlab code from http://yarpiz.com/255/ypml110-dbscan-clustering). The algorithm first finds neighboring data points within a sphere of radius *r*, and adds them into same group. In parallel, a predefined threshold minimal points (*minPts*) was used by the algorithm to justify whether any counted group is a cluster. If the number of points within a group is less than the threshold *minPts*, the data point is classified as noise. As a negative control, we generated uniformly sampled data sets with the same localization density as our ATAC-PALM localizations. We then implemented DBSCAN analysis by using 150 nm as the searching radius (*r*) (peak radius from the Ripley’s H function analysis) and empirically setting *minPts* as 10. To reconstruct the iso-surface for each identified ACD, the convex hull which contains the ACD data points was calculated and visualized by using Matlab. The volume of the convex hull was computed, and the normalized cluster radius (calculated from a sphere with equal volume) was estimated and shown in violin plot.

### Oligopaint FISH experiment and analysis

The Oligopaint FISH probe libraries were constructed as described previously (Beliveau et al., 2012). A Tier 15 ssDNA oligo pool was ordered and synthesized from Twist Bioscience (San Francisco, CA). Each oligo consists of a 32 nucleotide (nt) homology to the mm9 genome assemble discovered by the OligoArray2.0 with the following parameters −n 22 −D 1000 −l 32 −L 32 −g 52 −t 75 −T 85 −s 60 −x 60 −p 35 −P 80-m “GGGGG;CCCCC; TTTTT;AAAAA” run from the algorithm developed from the laboratory of Dr.Ting Wu (https://oligopaints.hms.harvard.edu/). Each library subpool consists of a unique sets of primer pairs for orthogonal PCR amplification and a 20 nt T7 promoter sequence for *in vitro* transcription and a 20 nt region for reverse transcription. Individual Oligopaint probes were generated by PCR amplification, *in vitro* transcription, and reverse transcription, in which ssDNA oligos conjugated with ATTO565 and ATTO647 fluorophores were introduced during the reverse transcription step as described previously (Beliveau et al., 2015; Boettiger et al., 2016). The Oligopaint covered genomic region (mm9) used in this study are listed below:

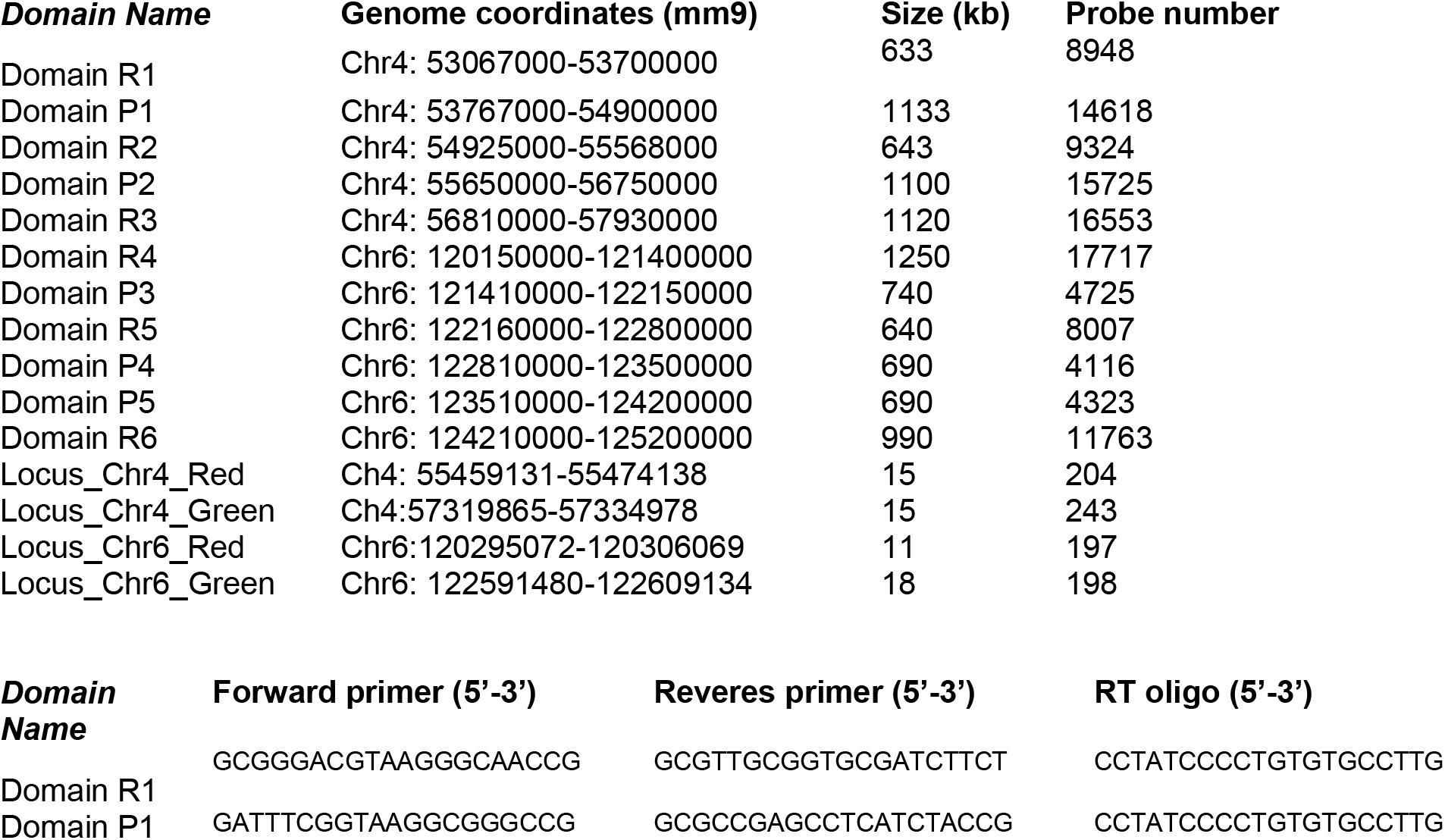
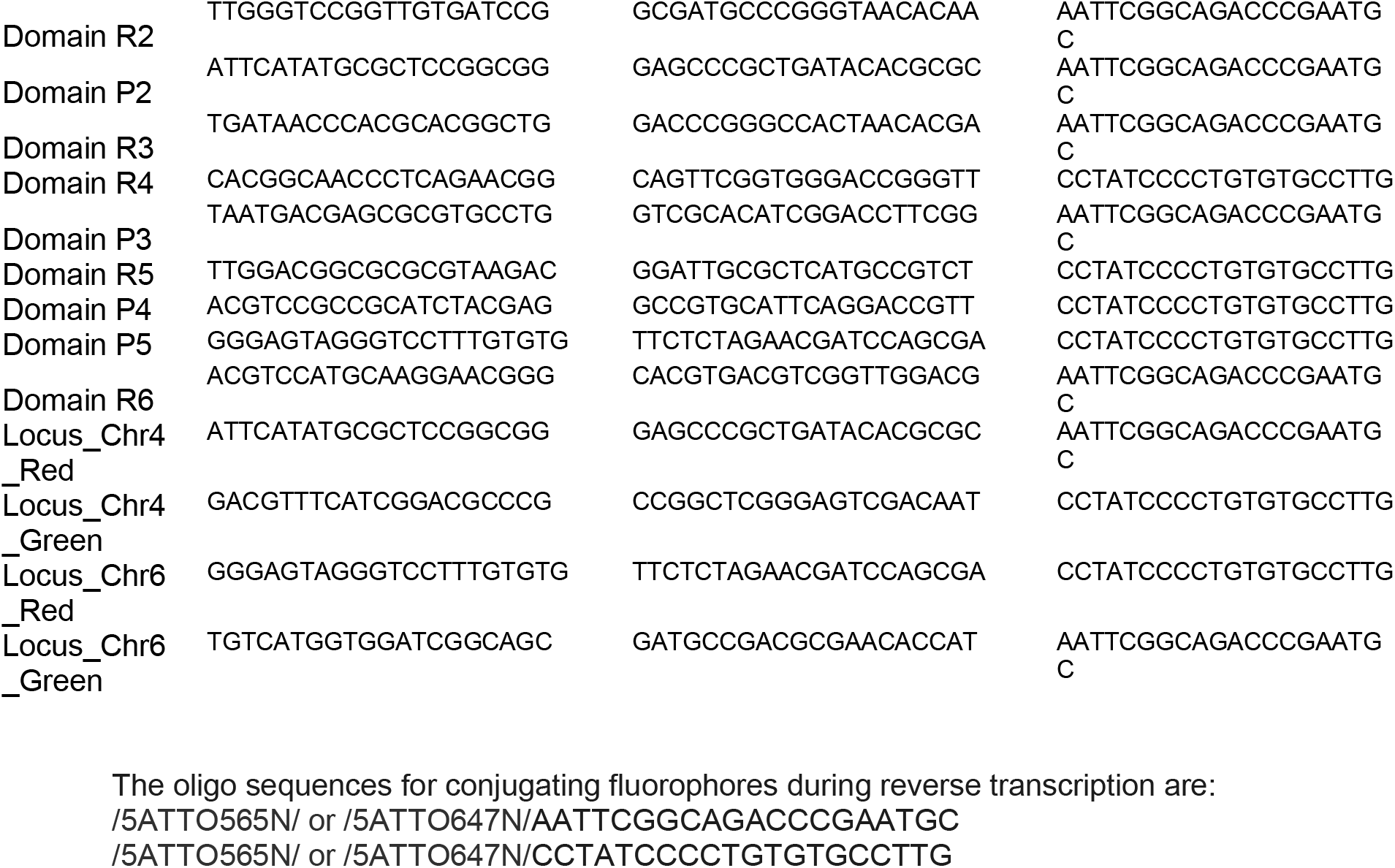

For 3D DNA FISH on ESCs, #1.5 round glass coverslips (Electron Microscopy Sciences) were pre-rinsed with anhydrous ethanol for 5min, air dried, and coated with 0.1% gelatin or equivalent for at least 2 hours. Fully dissociated ESCs were seeded onto the coverslips and recovered for at least 6 hours before experiments. Cells were fixed with 4% (v/v) methanol free paraformaldehyde (Electron Microscopy Sciences, Cat# 15710) diluted in 1X PBS at room temperature for 10min. Then cells were washed 2X with 1XPBS and permeabilized in 0.5% Triton-X100 in 1XPBS for 30min. After 2X wash in 1XPBS, cells were treated with 0.1M HCl for 5min, followed by 3X washes with 2XSSC and 30 min incubation in 2X SSC + 0.1% Tween20 (2XSSCT) + 50% (v/v) formamide (EMD Millipore, cat#S4117). For each sample, we prepare 25ul hybridization mixture containing 2XSSCT+ 50% formamide +10% Dextran sulfate (EMD Millipore, cat#S4030) supplemented with 0.5μl 10mg/mL RNaseA (Thermo Fisher Scientific, cat# 12091-021) +0.5μl 10mg/mL salmon sperm DNA (Thermo Fisher Scientific, cat# 15632011) and 20pmol probes with distinct fluorophores. The probe mixture was thoroughly mixed by vortexing, and briefly microcentrifuged. The hybridization mix was transferred directly onto the coverslip which was inverted facing a clean slide. The coverslip was sealed onto the slide by adding a layer of rubber cement (Fixo gum, Marabu) around the edges. Each slide was denatured at 78°C for 3 min followed by transferring to a humidified hybridization chamber and incubated at 42°C for 16 hours in a heated incubator. After hybridization, samples were washed 2X for 15 minutes in pre-warmed 2XSSCT at 60 °C and then were further incubated at 2XSSCT for 10min at RT, at 0.2XSSC for 10min at RT, at 1XPBS for 2X5min with DNA counterstaining with DAPI. Then coverslips were mounted on slides with Prolong Diamond Antifade Mountant (Thermo Fisher Scientific Cat#P36961) for imaging acquisition.

3D DNA FISH images were acquired on the ZEISS LSM 880 Inverted Confocal microscope attached with a Airyscan 32 GaAsP (gallium arsenide phosphide)-PMT area detector (Huff, 2015). Before imaging, the beam position was calibrated centering on the 32 detector array. Images were taken under the Airyscan Super-resolution mode with a Plan Apochromat 63X/NA1.40 oil objective in a lens immersion medium having a refractive index 1.515. We used 405nm (Excitation wavelength) and 460nm (Emission wavelength) for the DAPI channel, 561nm (Excitation wavelength) and 579nm (Emission wavelength) for the ATTO565 channel and 633nm (Excitation wavelength) and 654nm (Emission wavelength) for the ATTO647 channel. Z-stacks were acquired under Super-resolution mode for the optimal z sectioning thickness around 190nm. The Airyscan super-resolution technology used a very small pinhole (0.2AU) at each of its 32 detector elements to increase SNR ~4-8 fold and enables ~1.7-fold improvement of resolution upon linear deconvolution in both lateral (xy) and axial (z) directions. After image acquisition, Airyscan image was post-processed and reconstructed using the provided algorithm from ZEISS LSM880 platform.

3D DNA FISH analysis was performed in Imaris 9.1 installed in Windows 10 X64 OS with the GeForce GTX 760/PCIe/SSE2 (version 4.5.0 NVIDIA 369.09). We applied a background subtraction filter to the 3D Airyscan processed images before the downstream analysis. To characterize the 3D DNA FISH domain, we employed the synthetic model—Surfaces object from Imaris and applied a Gaussian filter (σ = 1 voxel in xy) before the downstream 3D segmentation and quantification. The 3D volume of the DNA FISH defined domain is estimated by the number of voxels within the detected objects with the voxel size (48.9nmX48.9nmX199nm).The sphericity score is estimated by the ratio of the surface area of a sphere to that of the object (with equal volume) defined by equation 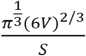 (V and S are volume and surface area of the object, respectively). To measure the 3D distance between foci-pairs, we localized the voxels corresponding to the local maximum of identified DNA FISH signal using the Imaris spots function module and calculated the Euclidean distance accordingly. To account for the chromatic aberrations between two channels in the imaging platform, we imaged the 0.5 μm multi-spectral beads (Zeiss) under the same acquisition conditions and calculated the distance difference between the channels of the same beads. And this distance was subtracted from all the downstream 3D loci-pair distance measurements.

For two-color 3D Oligopaint FISH and ATAC-PALM imaging experiments, cells were first mounted onto the 5mm coverslip embedded with nano-gold fiducial and treated with Tn5-PA549 transposome. With minimal exposure to light, samples were further processed for 3D DNA FISH experiments, except that HCl treatment is omitted and replaced with serial dehydration (immersed in 70%, 80%, 95% and 100% ethanol for progressive dehydration) step to maximally preserve the ATAC-PALM signal during imaging. The image stack of Oligopaint FISH channel was taken before the ATAC-PALM experiment under the Lattice light-sheet microscopy. The Oligopaint FISH image stacks were de-skewed and de-convoluted. The centroid of the diffraction limit spot of DNA FISH signal was identified by 3D Gaussian fitting and a cluster of 200 points were generated centering on the centroid to visualize the Oligopaint FISH signal represented in ViSP through Gaussian random sampling with a standard deviation σ=200 nm.

### Single particle tracking experiment and analysis

Single particle tracking (SPT) experiments were primarily carried out as described in (Chen et al., 2014b; Xie et al., 2017). Briefly, cells were seeded on 25mm #1.5 coverglass pre-cleaned with KOH and ethanol and coated with Matrigel according to manufacturer’s instruction. All live cell imaging experiments were conducted using an ESC imaging medium composed of FluoroBrite DMEM (Thermo Fisher Scientific, cat# A1896701) plus 15% FBS, 1XGlutaMax, 1XNEAA, 0.1mM 2-mercaptoethanol and LIF.

To analyze fast TF dynamics, 5 nM PA-JF_549_ HaloTag ligand were added to cells for 20 min and then cells were washed with imaging medium for 3 times for 15 min inside the 37°C incubator. Stroboscopic single particle tracking was performed using a custom-built microscope equipped with an Olympus 60× NA 1.49 TIRF objective and a custom tube lense (LAO-300.0, Melles Griot), resulting in 100x overall magnification as described in (English and Singer, 2015). An iXon Ultra EMCCD camera (DU-897-CS0-BV, cooled to −80 °C, 17 MHz EM amplifiers, preamp setting 3, Gain 400, ROI-height: 35 pixels) was synchronized using a National Instruments DAQ board (NI-DAQ-USB-6363) at a frame time of 5 ms. 2 ms stroboscopic excitations of a 555 nm laser (CL555-1000-O with TTL modulation, CrystaLaser) was synchronized to the frame times of the camera via LabVIEW 2012 (National Instruments). The laser stroboscopically illuminated the sample using peak power densities of ~1.7 kW/cm^2^ using HiLO illumination of the nucleus. We adjusted the TIRF illuminator to the HiLO mode to deliver a highly inclined illumination to the cover glass with an incident angle smaller than the critical angle for TIRF. The resulting laser beam forms a thin laminated optical sheet (~3 μm in thickness) that causes lower out-of-focus excitation, and thus higher SNR compared to conventional Epi-illumination (Tokunaga et al., 2008). PA-JF_549_ labels were photoactivated by 100-μs-long excitation pulses of 407 nm light (50 W/cm2) every 50 ms. During the course of image acquisition, the pulse interval was shortened to 25 ms. 10,000 frames were typically recorded per cell. During imaging, cells were maintained at 37 °C and 5% CO_2_ using a Tokai-hit stage top incubator and objective heater.

To study the TF stable binding events, cells labeled with JF_549_ HaloTag ligand were mounted onto a high speed motorized Nikon Eclipse Ti-E inverted microscope equipped with 100 x Apo TIRF 1.49 NA objective with correction collar, four laser lines (405/488/561/642 nm) and TIRF quad cube (405/488/561/640nm reflection bands), an automatic TIRF illuminator with motorized X axis and manual Y axis for beam positioning and focus, perfect focus 3 system, Triple DU-897 iXon Ultra EMCCD cameras on a Cairn Tri-cam emission splitter with filters 525/50 (GFP) 600/50 (RFP) and 705/72 (Cy5), a humidified incubation chamber maintained at 37°C with 5% CO_2_ (Tokai Hit). SPT was performed around 50 W cm^−2^ and 500 ms acquisition time for around 500 frames. The excitation laser was controlled by an AOTF (acousto-optic tunable filter) and reflected into the objective by a multi-band dichroic (405/488/561/633 BrightLine quad-band bandpass filter, Semrock). The emission light was filtered by a single band filter centered at 593 nm placed in front of the camera. To minimize drift during imaging, the environment control chamber was fully pre-thermo equilibrated and all imaging was conducted in an isolated ultra clean room with minimal mechanical vibrations. The Nikon NIS-Elements software was employed to control the microscope, laser lines and camera integration. We performed at least three independent biological replicates each with around 8 cells. Cells with obvious movement during acquisition were removed from further analysis.

We adopted a previously described analytical method—multiple target tracing (MTT) for SMT analysis (Sergé et al., 2008). Briefly, single fluorescent emitters were detected using a stringent generalized likelihood ratio test with very low false discovery rate. Detected peaks were evaluated by a 2D Gaussian fit with the enabled deflation loops. Localized particles were connected to generate trajectories by thresholding the diffusion coefficient by taking past history into account under maximum likelihood test. Blinking and photobleaching were also considered during tracking process. The single molecule tracking parameters are described previously (Xie et al., 2017).

TF SOX2 dynamics is thought to alternate between 3D diffusion and chromatin binding (specific or non-specific) according to our previous study (Chen et al., 2014). We measured the chromatin dissociation rate *k_m_* from 500ms acquisition time, wherein fast-moving molecules were motion blurred into background and escaped detection by MTT algorithm. To estimate *k_m_*, the maximum dissociation coefficient D was set 0.1 μm^2^ /s (estimated from H2B HaloTag) which constrains the identification of bound molecules for at least two consecutive frames. The chromatin residence time of bound molecules can be estimated from the dwell time calculated based on the trajectory length. Cells with apparent movement during imaging acquisition were removed from analysis. We generated the survival curve based on the 1-cumulative distribution function (1-CDF) histogram and mathematical fitting to resolve the dwell time of SOX2. Consistent with previous report, the one component decay model failed to the SOX2 trajectory survival curve. Instead, the dwell time of SOX2 is well fitted with a two-component exponential decay model described in Equation S17.

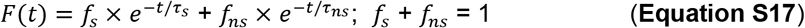

Two components are long-lived (*τ_s_*) and short-lived dwell time (*τ_ns_*) and their respective fractions (*f_s_* and *f_ns_*). *τ_ns_* could represent non-specific binding molecules or slow diffusion molecules. Because *τ_s_* largely disappear from tracking the SOX2 mutant lacking the DNA binding domain, the long-lived dwell time likely represents specific chromatin binding events in live cells (Chen et al., 2014b). Because molecules disappear from detection plane primarily by chromatin dissociation or photo-bleaching (after drift correction), the measured dissociation rate *k_m_* is described as the sum of true chromatin dissociation rate (*k_off_*) and photobleaching rate (*k*_b_)

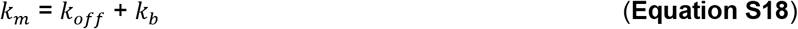

*k_b_* was derived through fitting the H2B HaloTag-JF_549_ survival curve, which accounts for the summed effect of photobleaching and drift (cells with apparent lateral drift were removed from downstream analysis).

To analyze SOX2 chromatin association kinetics, we performed stroboscopic illumination of SOX2 molecules at 500Hz with a camera integration rate 200Hz. The upper limit of diffusion coefficient for fast tracking was set 20 μm^2^/s and the jump length distribution, bound fraction, diffusion coefficient were analyzed by considering a steady-state TF two-state model (Hansen et al., 2017; Mazza et al., 2012). The model accounts for localization errors and corrects for freely-diffusing molecules gradually moving out-of-focus and we adopted a previously described MATLAB implementation of the model (Hansen et al., 2017, 2018).

Briefly, TF is assumed alternating between diffusion and chromatin bound state (S, substrate; C, TF-DNA complex):

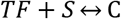

At steady-state, the bound fraction is estimated by

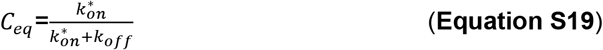

The average duration of two binding events for a given TF is estimated asStatistical Analysis

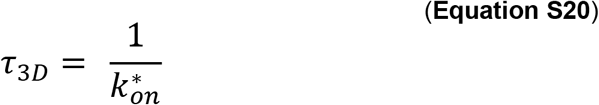

### Statistical Analysis

Unless stated specifically, data are presented as Mean ± SD (standard deviation) with statistical significance (* p < 0.05, ** p < 0.01, *** p <0.001). We used non-parametric Mann-Whitney *U* test to compare auto-correlation functions for 3D ATAC-PALM data. For other experiments involving relatively large number of measurements (Oligopaint FISH 3D distance, domain 3D voxels, TF dwell time and search time et al.), we employed two-tailed Student’s t-test.

### Polymer Modeling

We introduce a polymer model to study chromatin structure and dynamics. It explicitly considers the effect of Cohesin and TFs on chromatin organization. Since extrusion occurs on a much slower timescale than chromatin conformational rearrangement, we used a two-step algorithm to perform dynamical simulations. First, we carried out a one-dimensional (1D) stochastic simulation of extrusion to collect a set of Cohesin positions along the chromatin. Next, for each set of Cohesin configurations, we performed equilibrium molecular dynamics simulations to sample three-dimensional (3D) conformation of chromatin and chromatin bound protein molecules.

In the following, we provide details on the model setup and its dynamical simulation.

### 1D simulation of Cohesin extrusion

To determine the genomic locations of Cohesin molecules along the chromatin, we performed stochastic simulations of the extrusion model (Fudenberg et al., 2016; Sanborn et al., 2015) using the Gillespie algorithm (Gillespie, 1977). Two independent simulations were performed for the wild-type and CTCF depletion system, respectively. For each simulation, we recorded Cohesin positions along the genome at every time interval *τ*. The first 4-million-*τ* long segments of these trajectories were discarded as equilibration. Following the equilibration period, 100 sets of Cohesin positions were collected.

In these simulations, chromatin was modeled as a 1D lattice. Each lattice site corresponds to a genomic segment of 5kb in length, and a total of 600 sites is used to model a 3 Mb long segment. We further partitioned the chromatin into three regions of equal length, with the middle one as an ATAC-poor segment, and the two at the ends as ATAC-rich segments. ATAC-rich segments correspond to open chromatin and contain CTCF-binding sites, whose locations are detailed in the *Section: Detailed setup of simulation systems.* For simplicity, we assume the ATAC-poor segment does not contain open chromatin and is therefore free of CTCF sites.

The set of chemical reactions in this 1D model includes Cohesin binding, unbinding and extrusion (Fudenberg et al., 2016; Goloborodko et al., 2016). Cohesin molecules are modelled as two linked subunits, each of which occupies a lattice site. Cohesin molecules can bind onto the chromatin at a single lattice site or two adjacent sites with a rate of *k*_on_, provided that neither site is occupied by another Cohesin or CTCF molecule. The two binding modes ensures the formation of both odd and even sized loops. A constant rate *k*_on_ is used with the assumption that the nuclear protein concentration is high and remains constant. Once bound, Cohesin molecules can extrude to neighboring empty sites by moving both subunits in opposite directions at a rate of *k*_extr_, provided that the target sites are not occupied by other Cohesin subunits or CTCF. If the target sites are occupied by a CTCF molecule, the Cohesin is blocked but can move pass the CTCF molecule with a probability of 0.01, since these barriers are permeable as a result of protein unbinding. Bound Cohesin molecules can also dissociate from the chromatin at a rate of *k*_off_. Cohesin molecules were also assumed to fall off from the chromatin once they reach the two end beads.

To avoid boundary effects of the two end CTCF sites in each of the two ATAC-rich segments, we assumed that the first CTCF site does not block extrusion from the left side, and the last CTCF site does not block extrusion from the right side.

Numerical values for the rates used in the simulations are *k*_extr_ = *τ*^−1^, *k*_off_ = 0.04*τ*^−1^ and *k*_on_ = 0.002*τ*^−1^ respectively. From these rates, Cohesin processivity is estimated to be 250kb, and the mean separation between two neighboring Cohesin molecules is 90kb. These values are comparable to the optimal ones reported by Mirny and coworkers that were obtained by fine tuning the agreement between simulated and Hi-C contact maps (Fudenberg et al., 2016). Since extrusion rate has been estimated to be ~600bp/s (Terakawa et al., 2017; Ganji et al. 2018), the simulation time unit *τ* can be converted to real unit 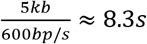.

### 3D simulation of chromatin organization

We carried out constant temperature and constant volume (NVT) simulations to model chromatin organization in 3D space using the software package LAMMPS (Plimpton and National, 1995). These simulations use the molecular dynamics algorithm to sample the equilibrium distribution of the following potential energy function that includes chromatin self-interaction, Cohesin mediated chromatin interaction, protein-protein, and protein-chromatin interaction

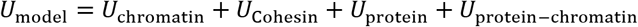

Detailed expression for each individual function is provided in the following using reduced units with length scale *σ* = 30 nm and energy scale *ϵ* = *k_B_T*. Temperature for all simulations were maintained at a constant *T* = 1.0 through Langevin dynamics with a damping coefficient of 0.5_τ_B__, where *τ*_B_ is the time unit. The time step is set to be đt = 0.01*τ*_B_. All simulations were carried out in a cubic box of length 60*σ* with periodic boundary conditions. Initial configurations of the chromatin polymer for all simulations are generated with a 3D random walk. Initial coordinates for protein molecules in the solution are also generated randomly.

The timescale in polymer models can be mapped onto physical units by matching the simulated diffusion constant with that in the nucleus. Specifically, the diffusion constant can be estimated using the fluctuation dissipation theorem as 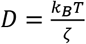. In Langevin dynamics, the friction coefficient 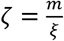, where 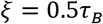 is the damping coefficient. The simulated diffusion constant therefore is 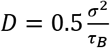, or 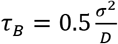. In the meantime, we can estimate the diffusion constant using the Stokes equation 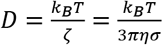, where *η* is the viscosity of the nucleoplasm, and has been reported to be on the order of 10 cP. Therefore, the time unit can be estimated as 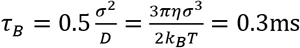.

#### Chromatin model and self-interaction

Chromatin is modeled as beads on a string. Consistent with the lattice model in 1D simulations, each bead is 5kb long, and a total of 600 beads was used to model a 3Mb long segment. The chromatin is again partitioned into three distinct segments, with the two end ones denoted as active (ATAC-rich segments) and enriched with CTCF and TF binding sites.

The potential energy function for the chromatin includes four terms,

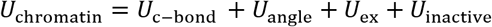

*U*_c-bond_(*r*) is the bonding potential between neighboring beads to ensure the connectivity and is defined as

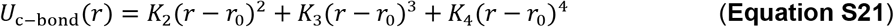

where 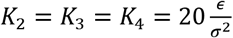 and *r*_0_ = 2.0*σ*.

*U*_angle_(*θ*) is the angular potential between every three consecutive beads to enforce the persistent length of the polymer. It adopts the following form

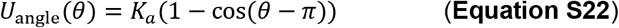

where *K_α_* = 2*ϵ*.

*U*_ex_ is the non-bonded potential to describe the excluded volume effect. It is applied between pairs of beads from ATAC-rich segments, and is modeled with the Weeks-Chandler-Andersen (WCA) potential

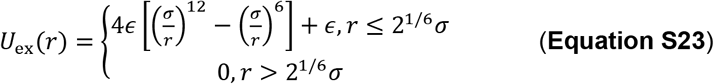

*U*_inctive_ is the non-bonded potential between beads from the ATAC-poor segment. It enforces a condensed conformation for the segment and adopts the form of a truncated and shifted Lennard-Jones potential

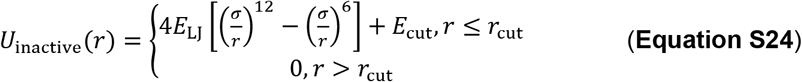

where *E*_LJ_ = 0.62*õ* and *r*_cut_ = 2.5*σ*. 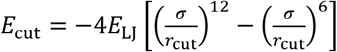 and ensures energy continuation at the distance *r*_cut_.

#### Cohesin mediated chromatin interaction

The effect of Cohesin extrusion on 3D chromatin organization was modeled using covalent bonds between two chromatin segments bound by a Cohesin molecule. Specifically, for each pair of chromatin beads occupied by the same Cohesin molecule as determined in 1D extrusion simulations, a harmonic bond of the following form is applied between them

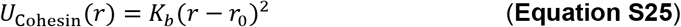

where *K_b_* = 4*õ* and *r*_0_ = *σ*.

#### Protein model and self-interaction

To account for the conformational flexibility of low complexity sequences in the TF activation domains and the multivalent interactions for TFs, we modeled TF molecules as tetramers with the two end monomers as binding domains. Binding domains interact favorably with specific protein binding sites on the chromatin, while non-binding domains show no such preference. In addition, a favorable interaction is introduced between binding domain themselves to promote protein-protein interaction. The energy function for protein is defined as

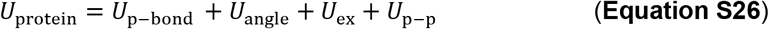

The bonding potential between neighboring protein monomers is defined as

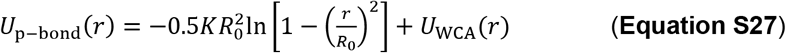

where 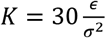 and *R*_0_ = 1.5*σ*. *U*_WCA_ is the same Weeks-Chandler-Andersen potential as that defined in Equation S23.

The angular potential *U*_angle_(*θ*) is the same as that for chromatin (Equation S22).

We used the same non-bonded potential *U*_ex_ defined in Equation S23 for excluded volume effect between pairs of non-binding beads and between non-binding and binding beads.

The final term *U*_p-p_ describes the favorable protein-protein interaction between binding domains and is modeled using the same truncated and shifted Lennard-Jones potential defined in Equation S24, except that the energy scale *E*_LJ_ = 2.5*õ*. We tested different values for the strength of protein-protein interactions, and found that under current model settings, *E*_LJ_ = 2*õ* is too weak to induce any chromatin clustering. On the other hand, *E*_LJ_ = 3*õ* is too strong and proteins will form clusters without the presence of chromatin. *E*_LJ_ = 2.5*õ* captures the effect that chromatin may act as nucleation sites for protein clustering.

#### Protein-Chromatin Interaction

The last term in the Hamiltonian, *U*_protein–chromatin_ describes the protein-chromatin interaction. The binding sites for TFs are located in the middle of two neighboring CTCF-binding sites, and a total of 15 sites are included in each ATAC-rich segment. For the specific interaction between protein binding domains and their binding sites on chromatin, a truncated and shifted Lennard-Jones potential defined in Equation S24 with *E*_LJ_ = 3*õ* is used. Otherwise, *U*_ex_ defined in Equation S23 is applied to model non-specific interactions between protein molecules and chromatin.

### Detailed setup of simulation systems

For 3D simulations, in addition to the chromatin, we introduced a total of 400 protein molecules into the simulation box. These proteins can exist either in an “active” or “inactive” state. Active protein molecules contain binding domains that can form specific interactions with chromatin. Inactive proteins that do not form specific interactions with chromatin serve mainly as crowding molecules that mimics the crowded environment of the nucleus. They also ensure that the impact of crowding on chromatin volume is similar in different setups when the concentration of active proteins is varied.

In extrusion-only, wild-type and CTCF-depletion systems, for each one of the 100 sets of Cohesin configurations obtained from the 1D simulations, we carried out a simulation that is 4 × 10^5^_*τ*_B__ in length. For the Cohesin-depletion system, a total of eight independent 10^6^_*τ*_B__-long simulations were performed. Configurations along the simulation trajectory were stored at every 2000 time-steps for analysis.

#### Extrusion-only model

As mentioned earlier, we explicitly considered the effect of CTCF molecules in hindering the movement of Cohesin along the chromatin during 1D simulations of extrusion. The two ATAC-rich segments harbor a total of 12 CTCF-binding sites that are uniformly distributed along the chromatin with a separation of 15 beads. CTCF sites start at the 15^th^ and 415^th^ bead for the two segments, respectively.

In accord with the original extrusion model, we did not consider the effect of TFs in 3D simulations. Therefore, all TF molecules are present in the inactive state.

#### Wild type

The locations of CTCF-binding sites are identical to those specified in the extrusion-only model. Out of these 400 molecules, only 75 of them are “active” and contain binding domains that can form specific interactions with chromatin. The concentration of active protein molecules is therefore 20.4 nmol/L, which is in the range of the reported nuclear protein concentration (Hancock, 2007).

#### CTCF depletion

In this system, CTCF sites are removed from the two ATAC-rich segments during 1D extrusion simulations. CTCF depletion will affect the average loop size formed by Cohesin molecules, which in turn affects the chromatin organization in 3D.

For 3D simulations, we kept the setup for TF molecules and their binding sites unchanged as those in the wild-type system. This is motivated by the observation that CTCF depletion has negligible effect on chromatin accessibility in the enhancers and promoters as shown in **Fig. S8**.

#### Cohesin depletion

For this setup, no 1D simulations were performed since loops cannot form without Cohesin molecules.

For 3D simulations, we increased the number of active protein molecules from 75 to 200 that corresponds to an increase of the concentration from 21.3to 56.9 nM. This increase of protein concentration is mainly motivated by the observation that the effect of 1,6-Hexanediol is much stronger in the Cohesin removal system than that in the wild type (see **Fig. 5A**). This suggests that protein-protein interaction is more significant in the Cohesin removal system.

Biologically, Cohesin molecules could impact the strength of protein-protein interactions, possibly via c-Myc mediated regulation. It has been shown that Cohesin exhibits strong and positive regulation of c-Myc expression (Rao et al., 2017; Rhodes et al., 2010), which could also regulate chromatin decompaction and nuclear architecture (Kieffer-Kwon et al., 2017). Interestingly, c-Myc has been reported to stimulate the phosphorylated state of low complexity domain containing proteins (Cowling and Cole, 2007). Therefore, it is possible that these is an increase of unphosphorylated proteins that can bind to chromatin and self-aggregate following Cohesin removal. It is worth noting that this Cohesin:c-Myc connection doesn’t preclude other uncharacterized mechanisms from contributing to the enhanced strength of protein-protein interactions upon Cohesin depletion.

#### 1,6-Hexanediol treatment

In this system, we adjusted the strength of specific interactions between protein binding domains (*U*_p−p_ in Equation S26) from 2.5*õ* to 2.4*õ*. This change is motivated by the role of the 1,6-Hexanediol in disrupting LCD-mediated protein-protein interactions. All the other simulation setups are kept the same as those in the wild-type system.

### Robustness of simulation parameters

A significant finding of the current study is that the extrusion model is insufficient to explain the enhanced clustering of accessible chromatin upon Cohesin removal. The rational for this is that loops enforced by Cohesin molecules tend to promote inter-chromatin contacts and favor collapsed chromatin conformations. Removing Cohesin in the extrusion model will always decrease chromatin compaction and, therefore, clustering. This conclusion should be general and independent from specifics of the model, as we show below.

#### Cohesin concentration

In order to demonstrate the robustness of this conclusion with respect to Cohesin concentration, we varied the binding rate *k*_on_ in 1D simulations from 0.002*τ*^−1^ to 0.004*τ*^−1^ and 0.001*τ*^−1^. These setups lead to normal, dense and light densities that correspond to a mean separation of 90kb, 180kb and 45kb between Cohesin molecules. As shown in **Fig. S10E**, for all the parameters explored, Cohesin removal always leads to a decreased clustering for chromatin.

#### Locations and density of CTCF-binding sites

In order to demonstrate the robustness of this conclusion with respect to CTCF locations, we performed additional simulations with an alternative distribution of the CTCF sites in each of the two active (ATAC-rich) segments. We kept the number of CTCF-binding sites the same but randomly generated an alternative set of CTCF sites on each of the active domains. In addition, we also performed simulations in which the total number of CTCF-binding sites on the chromatin was varied from 24 to 12 or 36. As shown in **Fig. S10F**, in all cases, removing Cohesin leads to a decreased clustering of accessible chromatin.

We further evaluated the robustness of simulation results from the polymer model that combines loop extrusion with the effect of TF molecules as follows.

#### CTCF positions

To test the robustness of results relative to CTCF locations, we performed additional simulations using the randomly generated CTCF sites mentioned above. As shown in fig. S12, D-G, the trends in chromatin clustering, 3D distance between active domains and volume of individual domains are preserved and qualitatively similar to those shown in Fig. 5C, fig. S11, C-D.

#### Protein concentrations

One assumption of the polymer model is the increase of the concentration for TFs upon Cohesin removal. To investigate the robustness of simulation results with respect to this concentration change, we tested a series of TF concentrations and compared all the metrics that we used: pair auto-correlation function, simulated FISH measurements and volume distributions of single ATAC-rich segments. (See **Fig. S12, A-C**). We found that beyond a certain threshold (56.9 nM, i.e., an increase by a factor of 2.7), all results are stable regardless of the protein concentration.

### Structural analysis of chromatin

#### Contact Probability map

We calculated the contact probability maps to highlight the contact difference between different perturbation conditions. We calculated the contact probability *P*(*r*) of each pair of beads with separation of *r* using the following expression (Zhang and Wolynes, 2015):

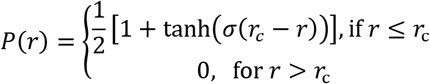

Where *r_c_* = 1.76 and *σ* = 3.72.

#### Pair auto-correlation function g(r)

*g(r)* was calculated as the probability distribution of pair-wise distances using the standard algorithm (Frenkel and Smit, 2001). We further normalize it with the corresponding distribution of hard spheres to highlight the clustering. The hard sphere *g(r)* was calculated from a uniform distribution for the same number of beads in a cubic box of the same size as the simulation box.

#### Volume for active domain

To calculate the volume of a given chromatin conformation, we determined the convex hull for all the 3D points that correspond to the Cartesian coordinates of chromatin monomers. The volume of the chromatin domain is then set as the volume of the convex hull.

### Analysis of transcription factor dynamics

To characterize the dynamics of transcription factor clusters, we measured the local protein density for the chromatin binding domains of each protein molecule using the following expression

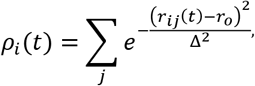

where *r_ij_*(*t*) is the distance between two chromatin binding domains *i* and *j* at time *t*, *r_o_* = 2*σ*, and Δ = 3*σ*. *j* iterates over all the chromatin binding domains of active proteins.

The time evolution of calculated local density profiles exhibits a two-state behavior as shown in **Fig. 6D, Fig. S14C**. These two states correspond to TFs moving in and out of clusters, respectively. To estimate the dwell time and search time of TFs, we then partitioned the trajectory into two states using a threshold of 0.5. For example, a TF will be classified as in a bound state if *ρ_i_*(*t*) ≥ 0.5; otherwise, the TF is unbound. We tested several threshold values, and found that all the results are robust as long as the threshold is below 1.0. We then estimated the dwell time as the mean time protein molecules remain in the bound state, and the search time *τ*_3D_ as the mean time protein molecules remain in the unbound state before finding their targets.

## Acknowledgements

We thank Xavier Darzacq, Shasha Chong, Thomas Graham, David McSwiggen, Claudia Cattoglio for critical reading of the manuscript, the Tjian-Darzacq lab members for helpful discussions and Prabuddha Sengupta and Herve Rouault for advice on data analysis. We also thank Deepika Walpita and Kathy Schaefer with assistance for FACS experiments, Damien Alcor for Airyscan Imaging and Melanie Radcliff for assistance.

